# Action potential waveforms are state-dependent

**DOI:** 10.64898/2026.09.15.751814

**Authors:** Blanca Martin-Burgos, Ashley Juavinett, Pamela D. Rivière, Ryan Hammonds, Bradley Voytek

## Abstract

Action potentials are brief electrical impulses that form the mechanistic basis for how neurons communicate. While it is well known that the shape of action potentials can differ across neurons, one fundamental assumption is that the complex voltage waveforms of action potentials within a given neuron are reducible to binary spikes. This assumption has constrained our conception of possible neural codes to those amenable to binary signaling, such as rate, temporal, and population codes. Here, we show that action potential waveform variability is not random, but is, instead, state dependent. To show this, we parameterize action potential waveforms in a set of very high temporal resolution (200 kHz) intracellular action potential recordings. We show that an action potential is not a digital ‘1’, but is instead a rich signal whose fine-scale features influence the shape and timing of the next action potential and whose waveform is systematically biased by input drive. We then show that intracellular action potential waveforms can vary as a function of the extracellular local field potential, but do so heterogeneously, as a function of the field potential amplitude and standard deviation. Our results have profound implications for systems and computational neuroscience, especially regarding the development of next-generation, biologically-inspired artificial neural networks that incorporate waveform dynamics. Non-binary action potentials point to a broader landscape of possible neural codes, whereby neurons communicate not just via binary spikes, but through their state-dependent waveform features.

## INTRODUCTION

Over 100 years ago, Lucas (1909) and Adrian (1914) performed seminal research demonstrating that the amplitude of a neuronal action potential (AP) is stable regardless of the intensity of the input stimulus. Rather than varying amplitude, neurons instead increase the number of APs that occur per unit time as a function of input intensity. These observations established the “All-or-none” Law, which states that an AP either occurs or does not; it is a binary signal whose amplitude is invariant to the inputs. This law fundamentally changed how we conceptualize computation in the brain. Under the assumption that this law is true, neural data analyses typically focus on identifying the rate at which APs occur or assessing the timing of these events in relation to other neural signals and behavioral states. This has constrained most theoretical models of neural coding to codes that are amenable to binary states such as rate codes, temporal codes, phase codes, and population codes (Ferster & Spruston, 1995; Gerstner et al., 1997; Maass, 1998; Ahmed & Mehta, 2009), all of which disregard the known variation in spike waveform shape between neurons. Interestingly, while the original neurons that composed early artificial neural networks (ANNs) assumed binary firing states (McCulloch & Pitts, 1943), modern ANN architectures implement continuous firing functions, which lead to superior performance (Nair & Hinton, 2010; Krizhevsky et al., 2012).

The reduction in complexity of AP waveforms to binary states is understandable from a historical perspective given the technical challenge of storing continuous electrophysiological time series sampled at 20 kHz or more. However, this signal reduction overlooks the fact that APs are fundamentally continuous, analog signals that are shaped by known biophysical mechanisms. We have known this since the early days of our field: Hodgkin and Huxley’s (1952) quantitative description of the AP established that spike generation is not a simple threshold event but the product of multiple time-varying ionic conductances with distinct kinetics that are variable and subject to biological noise. The advent of patch-clamp electrophysiology elucidated the dynamics of these signals by enabling direct, high-resolution recordings of membrane voltage and ionic currents (Neher & Sakmann, 1976; reviewed in Verkhratsky & Parpura, 2014). As recording methodologies and resolution improved further, they provided insight into how ion channel dynamics shape spike waveform characteristics (Toledo-Rodriguez et al., 2004; Bean, 2007).

With this improved resolution, it also became clear that distinct neuron types exhibit characteristic differences in AP shape. One of the most widely replicated findings is the categorization of neuronal subtypes based on AP width, differentiating between fast-spiking inhibitory interneurons that exhibit narrower spikes and excitatory pyramidal neurons that exhibit broader ones (McCormick et al., 1985; Connors & Gutnick, 1990; Wilson et al., 1994; Brumberg et al., 1996; Rao et al., 1999; Frank et al., 2001). While AP width has been a reliable marker for distinguishing inhibitory and excitatory neurons, early research showed that other features also carry cell-type information, including AP amplitude and measures of depolarization and repolarization kinetics, with features such as rise time and half-height width used in combination to classify neuronal subtypes (McCormick et al., 1985; Kyriazi et al., 1996; Erisir et al., 1999; Toledo-Rodriguez et al., 2004). More recent studies, using both intracellular and extracellular recordings, have expanded on this idea by combining multiple waveform features to classify a broader range of neuronal types, linking waveform features to functional properties (Bean, 2007; Trainito et al., 2019; Lee et al., 2023), gene expression profiles (Martini et al., 2023, Toledo-Rodriguez et al., 2004), physiological signals (Mosher et al., 2020), and neuronal morphology (Haynes et al., 2024; Moubarak et al., 2022).

The observation that AP waveforms differ systematically across neuron types laid the foundation for spike sorting methods, which use waveform shape to assign spikes recorded in extracellular recordings – a cornerstone of cognitive and systems neuroscience – to putative single units. These approaches rely on the assumption that each neuron generates a relatively consistent and distinguishable waveform such that clustering algorithms can separate overlapping activity from multiple nearby neurons and background noise (Lewicki, 1998). Building on this framework, a wide range of new algorithmic advances has improved accuracy, scalability, and compatibility with high-density and chronically implanted recording systems (Bestel et al., 2012; Su et al., 2013; Caro-Martin et al., 2018; Gibson et al., 2012; Pachitariu et al., 2016; Buccino et al., 2022; Pachitariu et al., 2024).

Substantial evidence from both intracellular and extracellular recordings, however, shows that AP waveforms can also vary systematically within a single neuron. A well-documented example is adaptation, in which APs broaden and firing rate decreases as a stimulus persists (Adrian & Zotterman, 1926; Aldrich et al., 1979; Jackson et al., 1991; Shao et al., 1999; Geiger & Jonas, 2000). In extracellular recordings, the first spike in a burst is typically narrower and higher in amplitude than subsequent spikes (Kandel & Spencer, 1961). Even more striking is recent evidence that human single neuron waveforms can change dramatically at seizure onset, such that clustering-based spike-sorting algorithms either assigned the altered waveforms to different neurons or discarded them entirely due to perceived instability (Merricks et al., 2021). These results suggest that neuronal AP waveforms are altered by the state of the networks in which they are embedded, raising the possibility that a single neuron could be artificially split into two or more putative neurons based on AP waveform differences within a single neuron, depending on the broader neuronal state.

While there is ample evidence at the cellular and molecular levels that AP waveforms of individual neurons are dynamically determined by ion channel function, synaptic input, and neuronal morphology, whether such variability extends to the systems level remains largely unexplored. Here, we address whether and to what degree within-neuron AP waveforms are affected by the network state. This question can be approached by examining mesoscale signals such as the local field potential (LFP), which is widely used in systems and cognitive neuroscience to track brain and network states including oscillations (Donoghue, J. P. et al., 1998; Donoghue, T. et al., 2022) and aperiodic dynamics (Gao et al., 2017; Donoghue, T. et al., 2020; Medel et al., 2023). The LFP reflects the aggregate of synaptic and intrinsic transmembrane currents within a local neuronal population (Buzsáki et al., 2012; Herreras, 2016), effectively capturing the peri-somatic ionic milieu. This signal varies systematically with neuromodulatory tone, behavioral engagement, and sensory input (Donoghue et al., 1998; Liu & Newsome, 2006; Mazzoni et al., 2008; Lee & Dan, 2012).

This body of evidence motivates a shift in how APs are conceptualized in systems and cognitive neuroscience (**Fig. 1**). To investigate the functional significance of within-neuron AP waveform variability and bridge cellular-level mechanisms of spike generation with systems-level theories of neural coding, we developed an open-source parameterization approach, *spikewise*, that quantifies fine-scale features of both extracellular and intracellular APs using an overcomplete set of waveform features, extending beyond the width and amplitude measures most commonly used in most prior work. To explore whether this variability reflects differences in ongoing network dynamics and brain states, we leveraged two open-source datasets (**Fig. 2E-F**): pvc-6 (Berg, 2014) and spe-1 (Marques-Smith et al., 2018, 2020). The pvc-6 dataset consists of intracellular recordings from two mouse visual cortical neurons (one SST+ interneuron and one unmarked cell) subjected to direct current stimulation, recorded *in vitro* under whole-cell current clamp at an unusually high temporal resolution (200,000 Hz), allowing us to quantify fine-grained changes in spike shape across input conditions. The spe-1 dataset consists of simultaneous intracellular and extracellular recordings from 43 neurons (40 analyzed: 35 putative pyramidal cells and 5 putative interneurons) in rat somatosensory cortex *in vivo*, with extracellular data from Neuropixels probes. Using pvc-6, we first show that AP features are systematically modulated by stimulation parameters. We then extend this analysis using spe-1 to examine whether changes in the peri-somatic LFP, a proxy for local network state, co-vary with spike waveform shape.

**Fig. 1.**
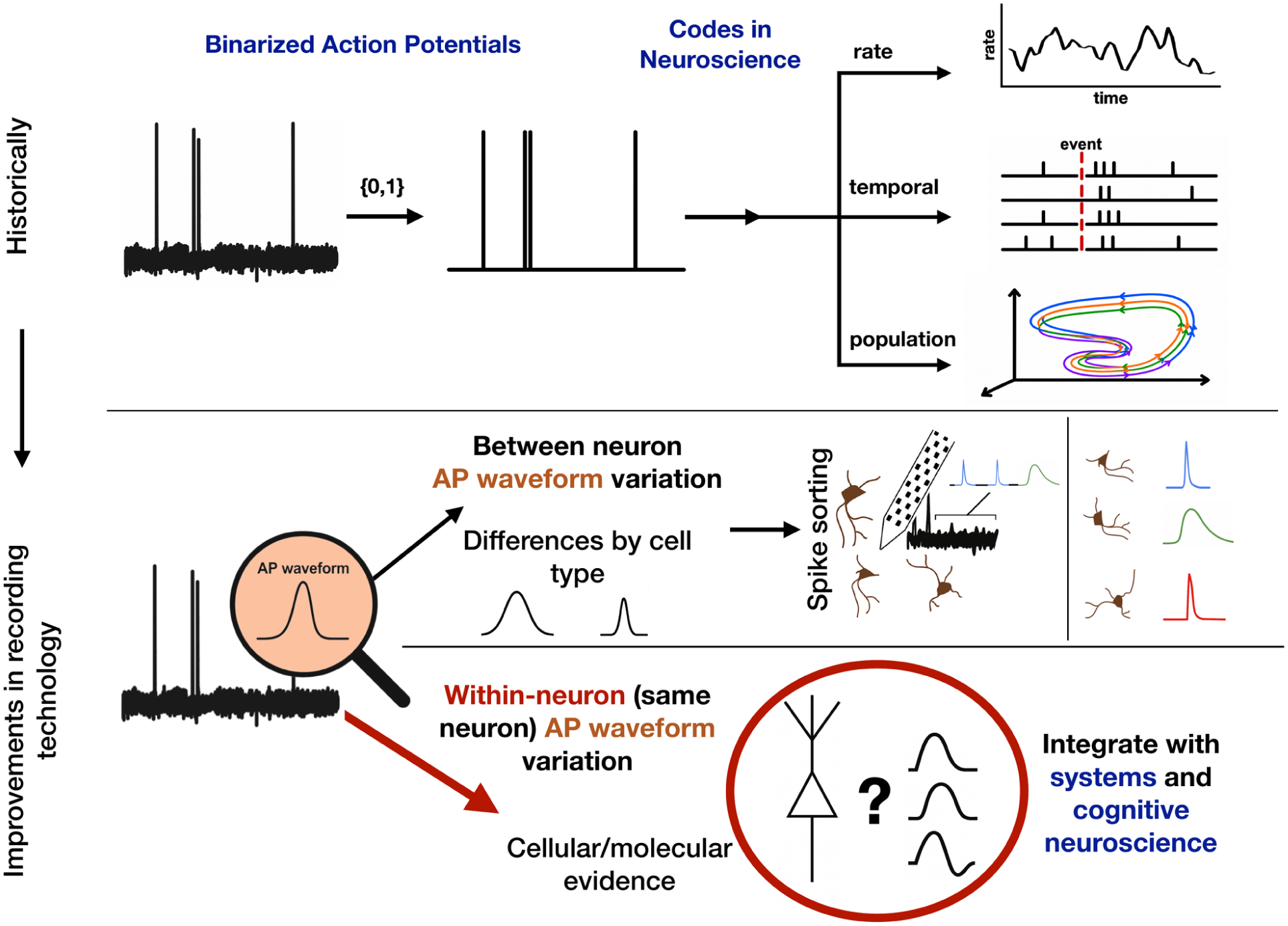
From binary spikes to dynamic waveforms. Within-neuron AP variability may contain meaningful information that is ignored in systems, cognitive, and computational neuroscience. (Top) Action potential waveforms are binarized and reduced to timestamps of when they occurred. These binary features are then used in much of computational, systems, and cognitive neuroscience to infer the “codes” that neurons used to process and / or transmit information. Common codes rely on counting how many spikes occur in some unit of time (rate coding), when the spike occurred relative to an external or intrinsic event (temporal coding), or examining the joint activity across a population of neurons, including which neurons are active, their relative rates, and their correlations (population coding). (Bottom) While AP waveforms are used for spike sorting extracellular recordings into putative individual neurons, those waveforms are usually then discarded. Here, we examine whether and to what degree within-neuron AP waveform variability is unstructured, or whether it is systematic and reflects neuronal state, inputs and surrounding population activity.

**Fig. 2.**
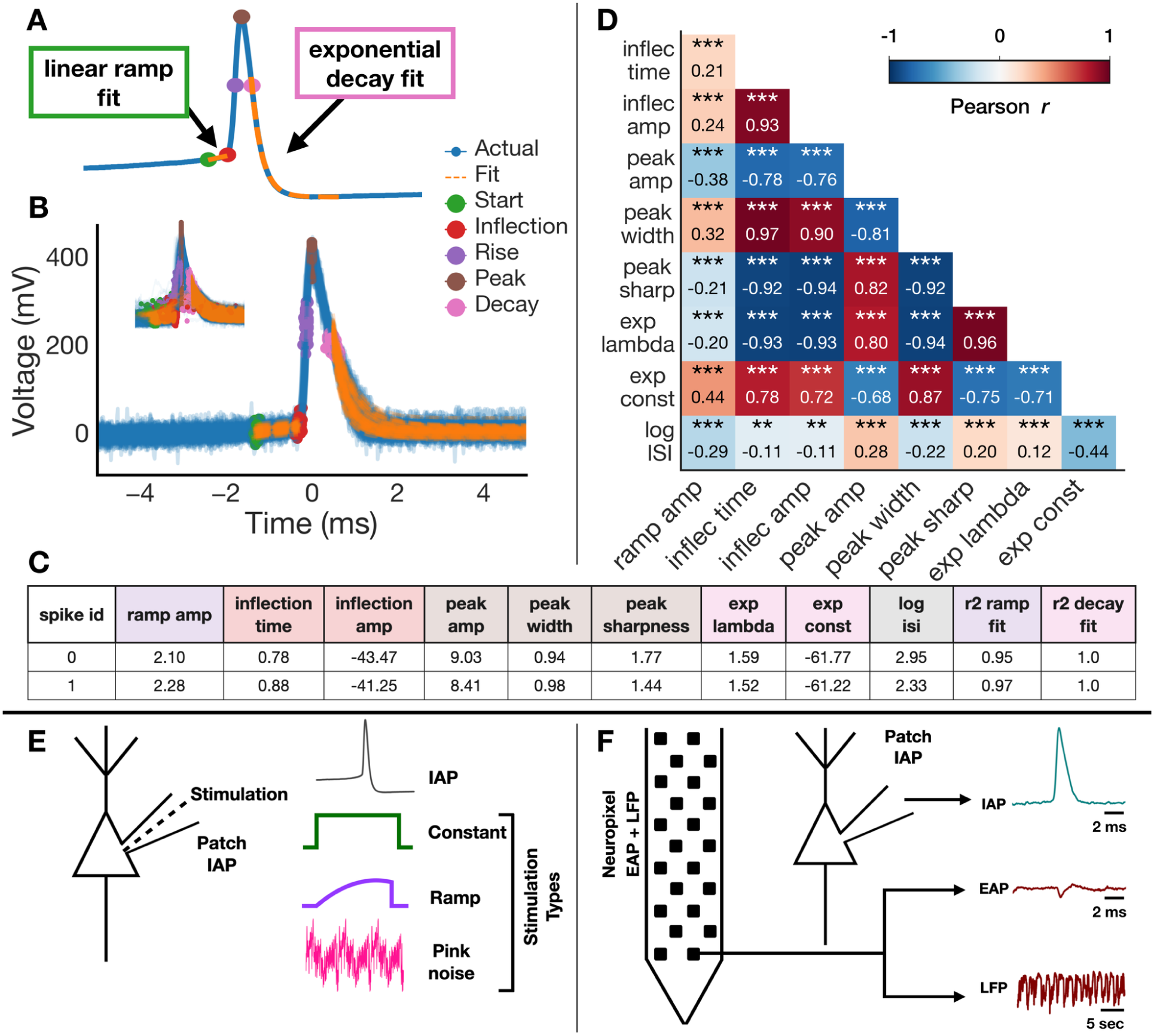
AP waveform parameterization and dataset schematics. **a**, Parameterization schematic: the pre-spike depolarization is fit with a linear ramp (slope = ramp amplitude; endpoint = inflection time and inflection amplitude); the peak is characterized by its amplitude, width (time between rising and decaying half-peak points), and sharpness (mean voltage difference between the peak and points 0.1 ms on either side); and the repolarization is fit with an exponential decay (decay rate λ, asymptotic constant). **b**, The *spikewise* parameterization method applied across action potentials recorded from a representative neuron (main plot: 100 representative spikes, inset: all spikes). **c**, Example dataframe of extracted AP waveform features where each row is a single AP parameterized by each column feature. **d**, Intra-neuron pairwise Pearson correlations reveal that individual spike features are systematically, and uniformly significantly, intercorrelated (data from pvc-6 Cell 1; SST+) (**p < 0.01; ***p < 0.001). **e**, Schematic of the Allen Institute (Berg, 2014) dataset, featuring high temporal resolution (200,000 Hz) intracellular recordings across varied stimulation protocols. **f**, Schematic of the spe-1 dataset (Marques-Smith et al., 2018), which includes simultaneous intracellular, extracellular, and LFP recordings.

## RESULTS

### Parameterization captures intra-spike correlations

Using our *spikewise* parameterization method on intracellular patch-clamp recordings of APs (**Fig. 2A**), we extracted a comprehensive set of waveform features, including ramp, inflection, and peak amplitudes; inflection time; peak width and sharpness; repolarization decay rate; and post-repolarization resting state amplitude. In addition, we computed the time to the next spike (inter-spike interval: ISI) and reported R² values for both the linear fit of the depolarization ramp and the exponential fit of the repolarization phase. From these extracted features, we observed significant within-neuron correlations (**Fig. 2D**).

Three relationships were consistent across all cells and both datasets: peak sharpness with peak width, peak width with inflection time, and peak sharpness with inflection time (**Supp. Fig. 1A-D**). Spikes with earlier inflection points tend to have wider, less sharp peaks. Other relationships varied across cells. In pvc-6 data, the exponential decay rate was strongly correlated with peak features (**Fig. 2D**). For example, individual APs with sharper peaks exhibited faster repolarization, and APs that decayed faster tended to be followed by shorter inter-spike intervals. These same patterns appeared in individual spe-1 cells, but the correlations attenuated to near zero when averaged across the population (**Supp. Fig. 1D**). The direction of the decay correlations differed between spe-1 cells (**Supp. Fig. 1E**): 26 were positive and 17 negative for decay rate against peak amplitude. Exponential fit quality did not predict the sign of this correlation (r = 0.03, p = 0.856), indicating that this heterogeneity may be biological rather than an artifact of poor fitting. Given the strength of these intra-spike correlations within individual neurons, it may be possible to predict aspects of AP shape, such as decay dynamics, based on earlier waveform segments like the ramp or inflection point, and use these dynamics to infer the timing of the next spike.

### Electrical stimulation causally influences spike waveform

To examine whether our parameterization methods captured changes in the waveform that result from inputs to the neurons, we used pvc-6 (**Fig. 2E**). First, we examined how a single neuron’s AP waveform varied as a function of stimulation type (constant, ramp, or pink noise) (**Fig. 3A**). We observed that the type of input significantly influenced the waveform structure. Using a random forest classifier trained on our extracted AP features, we were able to predict stimulation type with high performance (**Fig. 3B**: out-of-fold accuracy across 1000 repeated 5-fold splits: 78.4% ± 0.7%; chance: 33.3…%). This analysis was restricted to the cell with sufficient trials across all three conditions; the second cell had only one ramp sweep and was excluded (**Supp. Fig. 4**). Among the most informative features for classification were peak amplitude and repolarization dynamics (**Fig. 3C**).

**Fig. 3.**
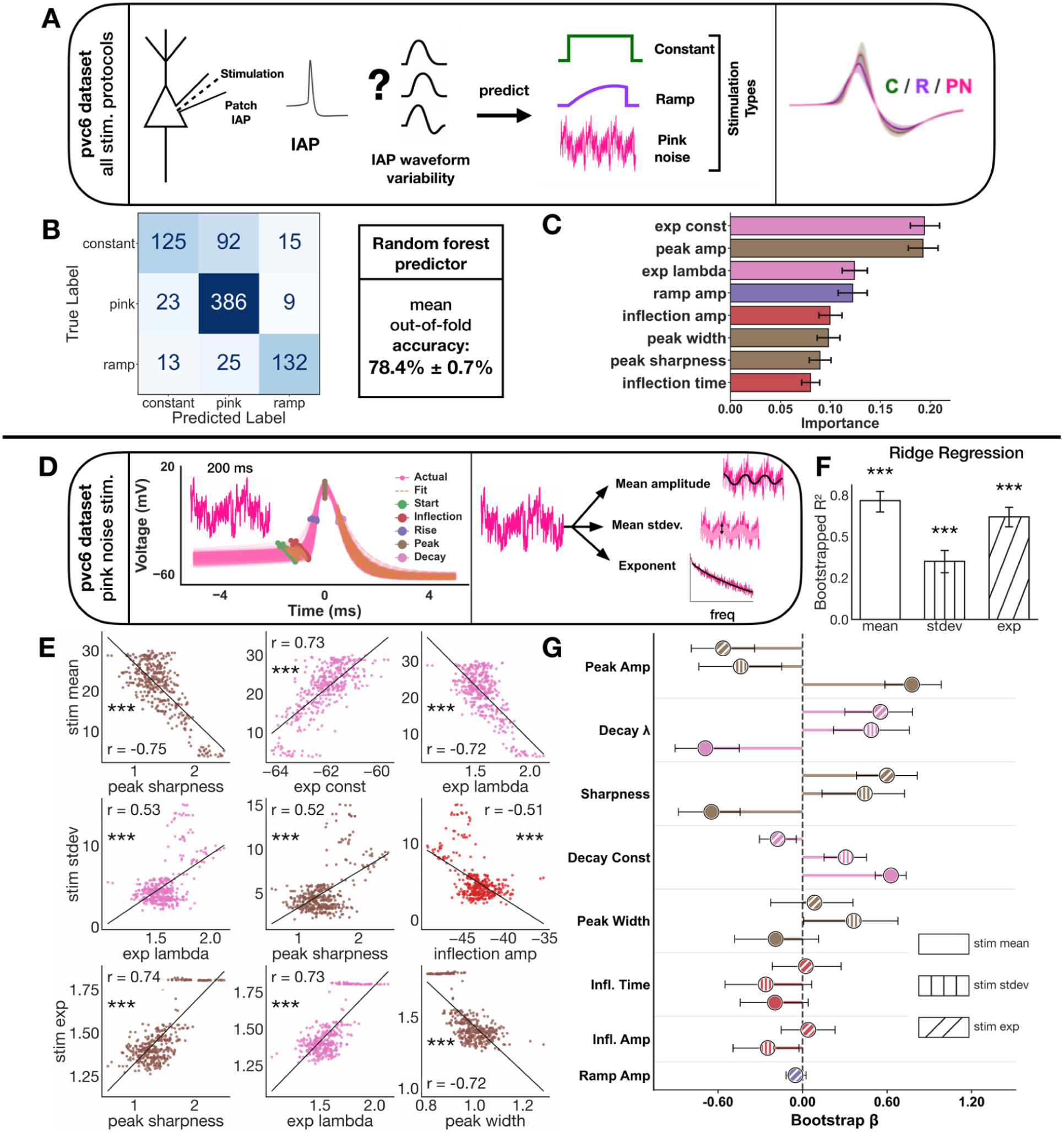
Electrical stimulation causally modulates AP waveforms. **a**, Dataset schematic: patch clamp intracellular AP recordings under three stimulation types (constant, ramp, pink noise); (right) mean waveforms overlaid by stimulation type. **b**, Confusion matrix for a random forest classifier predicting stimulation type from spike waveform features, averaged across 1000 independent out-of-fold 5-fold cross-validation splits so all 820 spikes are represented (hyperparameters selected by grid search; out-of-fold accuracy: 78.4% ± 0.7%). **c**, Random forest feature importances averaged across all fold-level estimators from the same 1000 repeated out-of-fold splits shown in b (error bars: 95% CI across folds). **d**, Ridge regression schematic: spike waveform features used to predict properties of pink noise stimuli extracted from a 200 ms window before each spike, including mean amplitude, standard deviation, and spectral exponent. **e**, Top pairwise Pearson correlations between spike waveform features and each pink noise stimulation target (significance: Pearson r, FDR corrected ***p < 0.001). **f**, Bootstrapped R^2^ for ridge regression models predicting each pink noise stimulation target from spike waveform features (alpha tuned via RidgeCV; model significance via permutation test, n=1000, FDR corrected ***p < 0.001). **g**, Significant bootstrapped beta weights (1000 iterations, 95% CI; one-sample t-test against zero, FDR corrected ***p < 0.001) per spike feature per pink noise stimulation target; dot fill encodes stimulus target (solid = stimulation mean, vertical lines = stimulation standard deviation, diagonal lines = stimulation spectral exponent).

We next examined the effects of pink noise stimulation, due to its dynamic fluctuations during recording and its spectral similarity to naturalistic 1/f-like neural activity, which may reflect fluctuations in brain state and underlying network dynamics (Gao et al. 2017; Medel et al., 2023). To characterize the pink noise stimulus, we computed the mean amplitude, standard deviation, and spectral exponent (Donoghue et al., 2020) of the injected current in a 200 ms window preceding each spike, and assessed how these properties related to AP waveform shape (**Fig. 3D**). Different spike waveform features were significantly correlated with all three stimulus properties (**Fig. 3E**). Using multivariate ridge regression, we found that AP waveform features significantly predicted all three stimulus properties in Cell 1 (SST+): current amplitude (R² = 0.72), amplitude standard deviation (R² = 0.35), and spectral exponent (R² = 0.62; all p < 0.001 by permutation test, FDR corrected; **Fig. 3F**). The dominant predictors differed by target: peak sharpness and exponential decay rate (λ) most strongly predicted mean current amplitude and spectral exponent, while peak width and decay offset drove standard deviation prediction (**Fig. 3G**). Expanding the pre-spike integration window revealed that prediction of mean amplitude continued to improve out to 500 ms before the spike occurred, while spectral exponent prediction rose steeply and then plateaued by 200 ms. Standard deviation prediction peaked at shorter windows (∼100 ms) and declined thereafter (**Supp. Fig. 5B**). A qualitatively similar but weaker pattern was observed in a second cell, with the best pre-spike windows differing across stimulus targets (**Supp. Fig. 5B-C**). These results suggest that stronger inputs not only increase neuronal firing rate, consistent with rate coding, but also modulate fine-scale features of a neuron’s AP shape. Waveform carried information about the mean amplitude, standard deviation, and spectral exponent of recent inputs, out to half a second before a spike. Across stimulation features, most of the dominant predictors involved the repolarization phase, pointing to a possible role for input-driven K⁺ channel dynamics.

### Neurons show multimodal within-cell spike waveform spontaneous variability

To test whether AP waveform also varies within neurons without experimenter-controlled input, we turned to the spe-1 dataset (**Fig. 2F**), which provides simultaneous intracellular AP, extracellular AP, and LFP recordings from the same cell (**Fig. 4A**). We parameterized the intracellular APs and observed that individual waveform features were often multimodally distributed within a single neuron. To quantify this, we clustered each feature independently within each cell (see **Methods**). Of 40 neurons analyzed, 30 (75%) exhibited significant multimodal clustering in at least one waveform feature, most often peak width and peak amplitude (14 cells each, 35.0%; **Fig. 4B**). The cluster-average waveforms were visually distinct in a subset of neurons (**Fig. 4C**; **Supp. Fig. 8**). Neurons also showed multimodal ISI distributions (85.0% of cells), but a spike’s waveform cluster did not predict whether it had a short or long ISI (median η² = 0.013; **Supp. Fig. 9**). Waveform state and firing rate therefore vary independently within a cell.

**Fig. 4.**
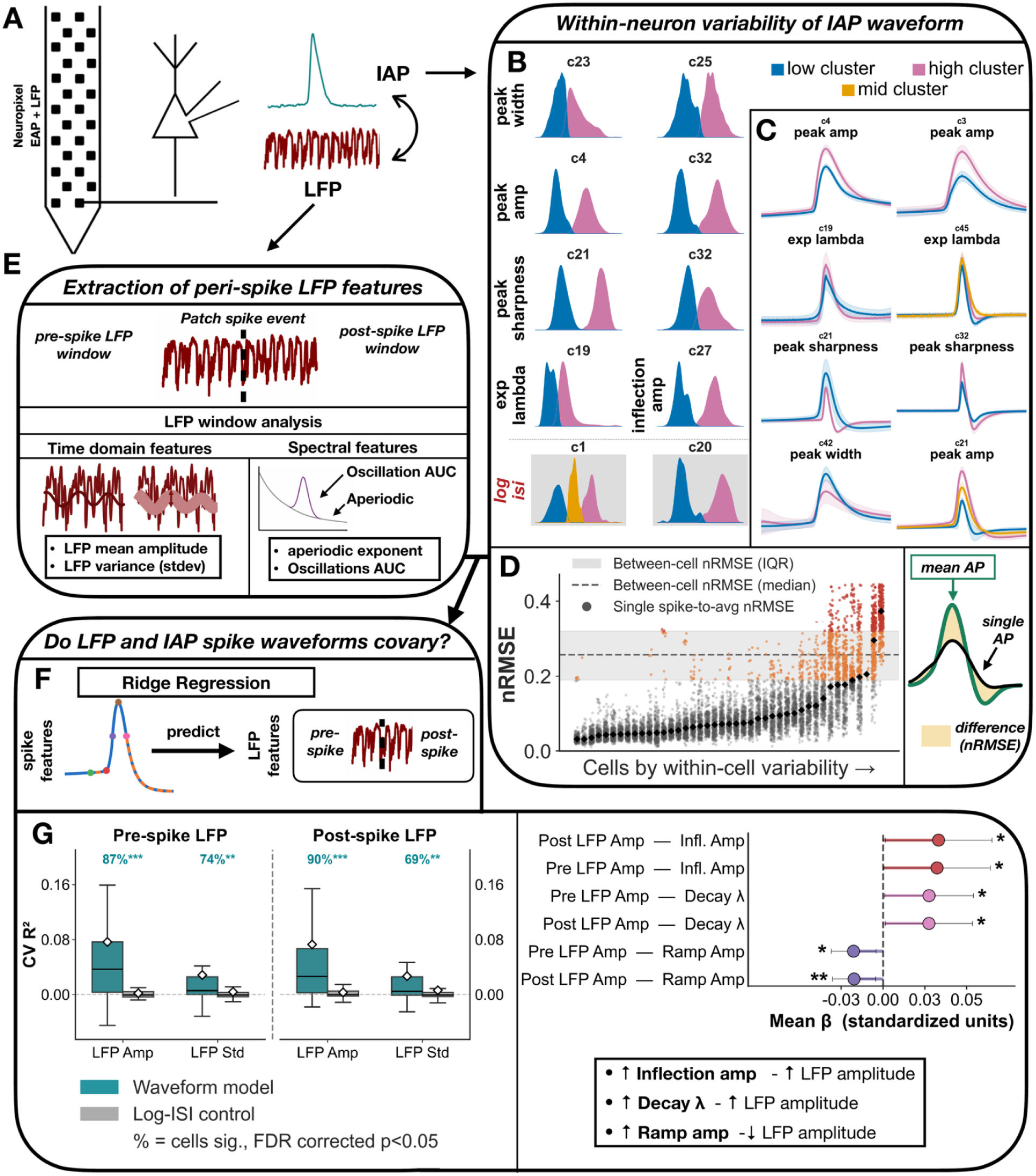
LFP state influences AP waveform within individual neurons. **a**, Dataset schematic: simultaneous whole-cell patch clamp intracellular AP, extracellular AP, and LFP recordings via Neuropixel probe. **b**, Within-neuron AP waveform variability across cells: spike feature distributions are multimodal in a subset of neurons, with example cells shown for peak width, peak amplitude, peak sharpness, exp λ, inflection amplitude, and log ISI (blue = low cluster, pink = high cluster, gold = mid cluster). **c**, Mean waveforms by cluster for representative cells, showing within-neuron AP waveform shape differences between clusters. **d**, Single spike-to-average nRMSE for all cells ordered by within-cell variability, showing that some individual neurons have higher waveform variability across APs within the same neuron than the average AP waveform variability between neurons (grey shading: between-cell IQR; dashed line: between-cell median; orange: high-variability cells); (right) schematic of the nRMSE metric. **e**, Peri-spike LFP feature extraction schematic: time domain features (mean amplitude, standard deviation) and spectral features (aperiodic exponent, aperiodic-adjusted oscillation power) extracted from pre-spike (−55 to −5 ms) and post-spike (+5 to +55 ms) LFP windows. **f**, Ridge regression schematic: spike waveform features used to predict pre- and post-spike LFP features. **g**, Left: CV R² distributions for waveform model versus log-ISI control across cells (% = fraction of cells significant, permutation test, FDR corrected p < 0.05; population Wilcoxon test, FDR corrected **p < 0.01; ***p < 0.001). Right: significant mean beta weights per spike feature per LFP target (one-sample t-test, *p < 0.05; **p < 0.01; standardized units).

These waveform states were not randomly distributed over the course of a recording. Correlating spike index with cluster label revealed significant temporal structure in most cell-by-feature pairs (88/103; **Supp. Fig. 10C**), such that a neuron might produce low peak amplitude spikes during one section of a recording and high peak amplitude spikes during another. In many cells these waveform state shifts were abrupt enough to localize in time: fitting a sigmoid to each cell’s rolling cluster trajectory identified discrete transition points between these states (**Supp. Fig. 11**). We tested whether recording metadata could predict the separation and temporal structure of these clusters using three measures: normalized root mean squared error (nRMSE; quantifies overall waveform difference including amplitude), cosine similarity (measures shape differences independently of amplitude), and temporal drift (reflects shifts in cluster membership over the recording; i.e., spikes changing from one feature group to another, such as narrow to wide spikes). No metadata predicted these measures (Mann-Whitney U, Kruskal-Wallis, and Spearman correlation with BH-FDR correction; all p > 0.05), nor which features exhibited multimodal clustering (chi-square; all p > 0.05).

### Within-neuron waveform variability can exceed between-neuron differences

We next compared how much waveforms vary within a neuron to how much they differ between neurons. Within-cell spike-to-average nRMSE was substantially lower than between-cell nRMSE (medians: 0.076 vs. 0.257; **Fig. 4D**; **Supp. Fig. 14A**), and the same was true for cosine similarity (0.984 vs. 0.762). In 2 neurons, however, the per-cell median nRMSE exceeded the between-cell median (**Fig. 4D**, rightmost cells; 97.6% and 66.5% of spikes above the between-cell median, respectively), and in 4 neurons more than 25% of spikes crossed that threshold. The same pattern held when spikes were compared only to cells of the same putative type or recorded with the same method (PC/PC, IN/IN, juxtacellular/juxtacellular, whole-cell/whole-cell; **Supp. Fig. 14B**). Across cells, the degree of separation was feature-dependent: ramp amplitude and inflection amplitude showed the most within/between overlap (within-cell median 49% and 62% of the between-cell median, respectively), whereas peak amplitude and peak sharpness had the clearest between-neuron separation (8% and 9%; **Supp. Fig. 14C**). Even when a cell’s whole-waveform median nRMSE fell below the between-cell level, individual spikes could exceed it: 8 cells had more than 10% of their spikes above the between-cell threshold (**Supp. Fig. 14D**). At the level of individual features this was far more widespread, with 32 cells each for ramp amplitude and inflection amplitude and 19 for exponential decay rate, whereas peak amplitude and peak sharpness remained near zero across all cells. Together, these results show that AP waveforms from the same neuron can sometimes look as different as, or more different than, waveforms from different neurons.

### AP waveform predicts peri-spike LFP state

We then used the LFP data from spe-1 to examine how network state might relate to AP waveform. For each spike we extracted LFP features from windows before (−55 to −5 ms) and after (+5 to +55 ms) it (**Fig. 4E**), capturing time-domain features (mean amplitude and standard deviation) and spectral features (aperiodic exponent, and oscillatory power in the theta and gamma bands, adjusted for aperiodic dynamics (Donoghue et al., 2022). (**Fig. 4E**). Spectral features were obtained by parameterizing the LFP power spectrum, which separates the aperiodic component from periodic oscillatory peaks so that oscillatory power is measured independently of broadband shifts (Donoghue et al., 2020). Within each of the 39 cells with usable LFP recordings, we used ridge regression to predict these LFP features from the eight waveform features. To test whether waveform shape predicts the LFP beyond what firing rate alone can, we ran the same models using log inter-spike interval as the only predictor (**Fig. 4F**). Pre- and post-spike beta coefficients were highly correlated across all features for LFP amplitude and standard deviation (Spearman ρ ≥ 0.89; **Supp. Fig. 15**), indicating that waveform shape related to the surrounding LFP similarly before and after the spike.

At the population level, cross-validated R² was significantly above zero for LFP amplitude and LFP standard deviation, but not for any of the spectral targets (Wilcoxon signed-rank on FDR-significant targets, **p < 0.01; ***p < 0.001; **Fig. 4G**, left; **Supp. Fig. 16A**). Waveform models also outperformed the log-ISI control for LFP amplitude (waveform: 87–90% of cells individually significant; ISI control: 46–51%) and LFP standard deviation (waveform: 69–74%; ISI control: 39–41%; permutation test, FDR corrected, p < 0.05). Prediction strength varied widely across cells, with mean CV R² well above the median for both targets (amplitude: 0.077 vs. 0.034; standard deviation: 0.030 vs. 0.006). Three waveform features carried consistent population-level weights for LFP amplitude: inflection amplitude (positive), decay rate (positive), and ramp amplitude (negative), indicating that spikes with a larger depolarizing inflection, faster exponential decay, and smaller ramp amplitude co-occur with higher-amplitude surrounding LFP (**Fig. 4G**, right). No individual feature reached consistent significance for LFP standard deviation, despite robust cell-level prediction. For six feature-target pairs, beta weights were large in individual cells but opposite in sign across them, resulting in no population-level effect (**Supp. Fig. 16C**). Although waveform feature contributions varied across cells, AP waveform shape overall better reflected surrounding population activity than spike rate alone.

## DISCUSSION

Models of neural coding in systems and cognitive neuroscience treat the AP as a binary event, while cellular electrophysiology has long documented that its shape varies with ion channel state, recent input, and firing history (Bean, 2007; de Polavieja et al., 2005; Shao et al., 1999). Whether this variability extends to the systems level, and whether it carries information about the state a neuron is embedded in, has remained largely unexamined. In this work, we parameterize AP waveforms from two open datasets and test these questions explicitly.

We observed reliable within-neuron waveform variability both under controlled stimulation and during spontaneous activity. Waveform features predicted the type and properties of injected current, and in the absence of controlled input, most neurons showed multimodal distributions of waveform features. In a subset of cells, the difference between individual spikes and their own cell’s average exceeded the typical difference between two different neurons. Spike sorting assumes the opposite: that each neuron produces a consistent and distinguishable waveform (Lewicki, 1998; Rey et al., 2015). If this assumption fails, a neuron whose waveform shifts with ongoing state, while its firing is otherwise unchanged, could be split across two sorted units. Each would carry part of the original spike train and appear to fire at a lower rate than the cell actually did, with implications for rate coding. In macaque prefrontal cortex, for example, sustained delay activity is taken as evidence that a stimulus is being held in working memory (Miller et al., 1996). A neuron firing steadily through such a delay, whose waveform varies across it, could be split into one unit active early and another active late, making it look stimulus-locked.

Beyond its consequences for spike sorting, waveform variation may offer insight into how neural activity shapes behavior, and vice versa. At the molecular level, modulation or deletion of specific potassium channels alters spike shape (Shao et al., 1999; Bean, 2007), and backpropagating AP amplitude determines the magnitude of the Ca²⁺ signal driving Hebbian plasticity (Magee and Johnston, 1997). These dynamic changes in ion channel function are influenced by neuromodulation (Cantrell and Catterall, 2001; Lee and Dan, 2012; Yang et al., 2013), synaptic input and recent membrane voltage history (de Polavieja et al., 2005), and axonal and dendritic morphology (Stuart et al., 1997; Moubarak et al., 2022). Moreover, AP waveform shape – not just its rate or timing – can directly affect downstream neuronal activity: increasing the width or duration of an AP at presynaptic terminals enhances both the amplitude and duration of excitatory postsynaptic currents in target neurons (Jackson et al., 1991; Sabatini & Regehr, 1997; Geiger and Jonas, 2000). These findings suggest that dynamic variation in spike waveform may serve as a mechanism for fine-tuned regulation of synaptic efficacy and, ultimately, behavioral output.

Waveform features also predicted the amplitude and standard deviation of the surrounding local field potential, beyond what firing rate alone explained. The LFP reflects synaptic and intrinsic transmembrane currents across the surrounding population (Buzsáki et al., 2012; Herreras, 2016), some of which may also shape a neuron’s own spike waveform. This relationship was modest and differed across cells: which features tracked the local field varied between neurons, so relationships that were strong within individual cells were weak once averaged across them. Given that AP waveform shape depends on membrane conductance in the milliseconds preceding spike initiation (de Polavieja et al., 2005), features encoded in the LFP could be mechanistically linked to spike shape, offering a bridge between intracellular dynamics and the large-scale signals typically studied in systems and cognitive neuroscience.

We considered several caveats and potential limitations in interpreting these results, though. Waveform variability in patch-clamp recordings has been taken as an indicator of a failing patch seal or a dying neuron, and recordings showing spike shape broadening and drift are routinely excluded from analysis on these grounds (Kandel, Spencer and Brinley, 1961; Ledochowitsch et al., 2019). If recording quality were driving our results, waveform parameter variability should increase over the course of a recording. Instead, the rolling interquartile range (IQR), which quantifies the spread of waveform parameter values over time, remained stable across normalized recording time. The distribution of temporal slopes was also centred on zero across all features and cells, with no positive skew (median 0.001; **Supp. Fig. 3**). We therefore find no evidence that recording quality contributed to the variability we report. Alternatively, our results could reflect adaptation or refractory-period dynamics rather than a distinct internal state. However, waveform clusters were largely independent of ISI-based clusters (median η² = 0.013; **Supp. Fig. 9**), and waveform features predicted LFP state beyond what log ISI alone explained (**Fig. 4G**), indicating that firing history alone does not explain the variability we report. We also cannot rule out that stimulation was delivered at some points during the spe-1 recordings to drive neuronal activity in low-firing neurons. The dataset is described as spontaneous activity recorded without experimenter-controlled stimulus, so this is unlikely to be a major factor, but the possibility remains. To test this possibility, we checked each cell for shifts in membrane potential, as this may be an indication of stimulation (**Supp. Fig 17**). A subset of cells showed baseline shifts that also coincided with waveform-shape shifts. Even so, these membrane potential shifts do not appear to be the sole driver of the AP waveform variability we report, since AP waveform shape continued to be variable in most of these cells after the shift, rather than switching to a single new state.

It is also important to note that all the results presented here are from intracellular data, where every recorded spike can be attributed to a single neuron and its waveform is set by that neuron’s own transmembrane currents. Extracellularly recorded waveforms vary for additional reasons: electrode position relative to the soma and dendrites strongly influences recorded shape (Gold et al., 2006), and probe drift alters this geometry over a session (Buccino et al., 2022). Measuring variability intracellularly therefore isolates the physiological component. The spike sorting implications we describe, however, apply to extracellular data. If anything, the differences we observe intracellularly would be expected to persist there, with recording-based variability added on top.

Several directions follow from these findings. First, it will be important to assess how specific waveform features vary within and between neurons and neuronal subtypes as a function of input, state, and individual unit physiology. Differences between cells were a recurring feature of our results: which features correlated with one another, which showed multimodal structure, and which tracked the local field all varied across neurons. Thus, characterizing this properly will require more cells, and datasets spanning more neurons would help establish which waveform relationships generalize and which are specific to individual cells. Second, further investigating the relationship between AP waveform variability and network dynamics or brain states, which our LFP analyses begin to address, could meaningfully extend conventional spike-based analyses and help link cellular mechanisms with systems-level processes, offering deeper insight into how neural activity supports cognition and behavior.

Together, these findings support studying within-neuron variability in AP waveform alongside traditional spiking metrics. They may also have implications for computational models of spiking activity. Spiking neural networks are built on discrete, binary spike events and encode information through rate, temporal, phase, or burst codes – the same schemes used to describe biological neuronal spiking. Such networks are valued both for consuming substantially less power than conventional artificial neural networks and for being more biologically realistic (Maass, 1998). Their biological realism, however, depends on the assumption that a spike is well described as a binary event, that is, it may be that neuroscience has mistaken the discreteness of spike occurrence for discreteness of spike function. If waveform shape carries information about recent input and network state, that means that binary units discard a dimension of the signal that they are meant to reproduce, potentially offering new paths toward more biologically realistic non-binary spiking neural network architectures.

## Data availability

All code used for all analyses and plots are publicly available on GitHub at https://github.com/voytekresearch/spikewise

## Acknowledgements

Support: e.g., NIH National Institute of General Medical Sciences grant R01GM134363-01 (to B.V.).

We thank the Allen Institute for Brain Science and André Marques-Smith, Adam R. Kampff, and colleagues for making the pvc-6 and spe-1 datasets publicly available through CRCNS.

## Author contributions

All authors conceived of the experiment(s) and developed the analyses, wrote analysis code, analyzed data, and wrote and edited the manuscript.

