## Supplemental Figures for "Action potential waveforms are state-dependent"

### SUPPLEMENTARY FIGURES

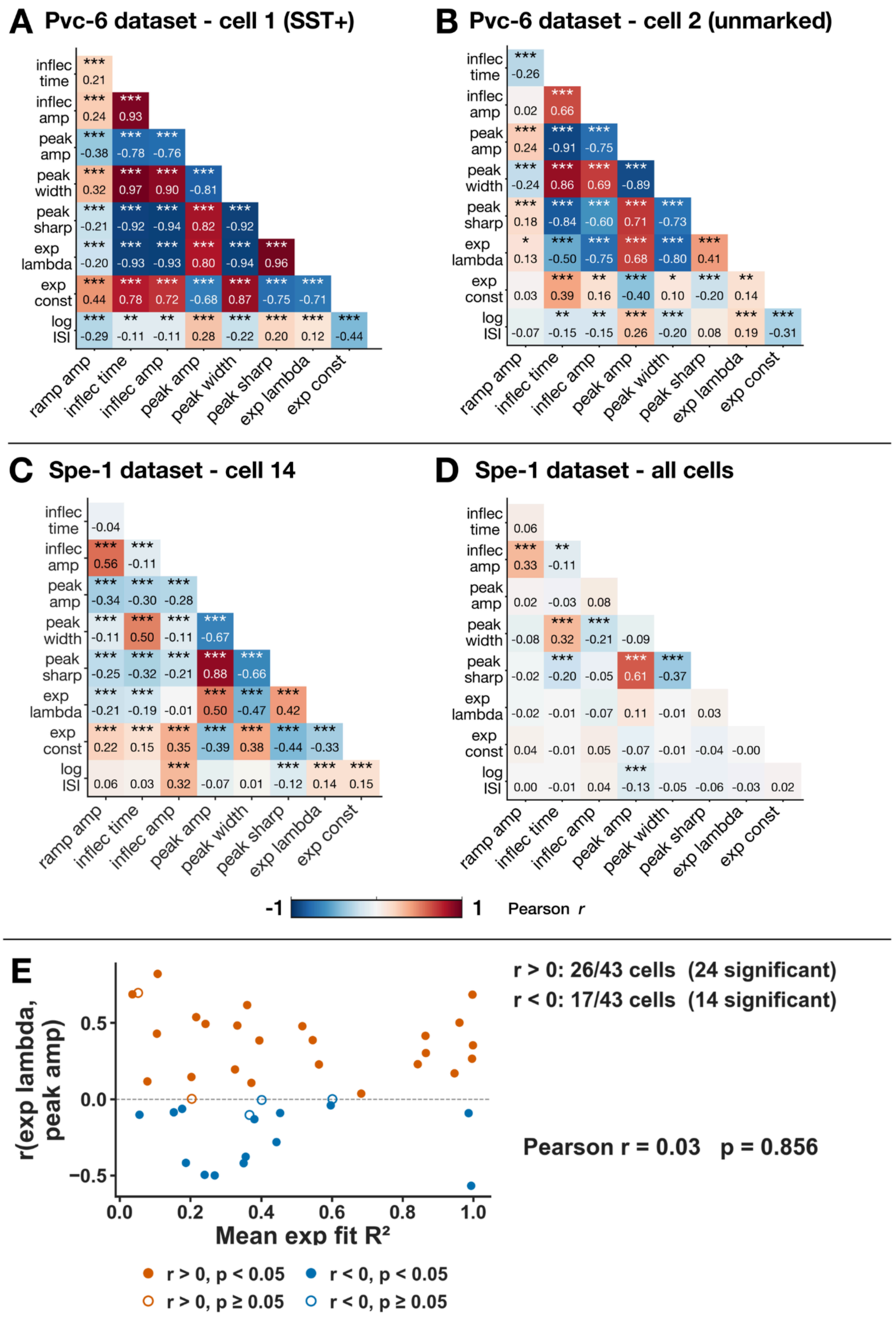

**Supp. Fig. 1 | Intra-spike waveform feature correlations across datasets and cell types.** Pairwise Pearson r between spike waveform features extracted by the parameterization algorithm (ramp amplitude, inflection time, inflection amplitude, peak amplitude, peak width, peak sharpness, exponential decay rate λ, exponential offset, log ISI). **a-c**, Lower-triangle heatmaps for individual cells; color indicates Pearson r (red, positive; blue, negative; scale −1 to 1); significance markers reflect raw pairwise Pearson r p-values (*, $p < 0.05$; **, $p < 0.01$; ***, $p < 0.001$). **d**, Mean within-cell Pearson r averaged across all spe-1 cells; significance reflects one-sample t-test against $r = 0$, FDR-corrected p (*$p < 0.05$; **$p < 0.01$; ***$p < 0.001$). **a**, pvc-6 Cell 1 (SST+; N = 820 spikes). **b**, pvc-6 Cell 2 (unmarked; N = 380 spikes). **c**, spe-1 Cell 14 (N = 6,101 spikes; best-matching spe-1 cell to pvc-6 Cell 1 by exp λ correlation sign profile, 7/7 matches). **d**, Mean within-cell Pearson r averaged across all spe-1 cells (N = 43 cells). **e**, Per-cell Pearson r between exponential decay rate (λ) and peak amplitude plotted against mean exponential decay fit quality ($R^2$) across all spe-1 cells with sufficient spikes (N = 43). Color indicates correlation direction (orange, $r > 0$; blue, $r < 0$); fill indicates significance (filled, $p < 0.05$; open, $p \geq 0.05$). Of 43 cells, 26 showed positive correlations (24 significant) and 17 showed negative correlations (14 significant). The absence of a relationship between fit quality and correlation direction (Pearson $r = 0.03$, $p = 0.856$) indicates that directional heterogeneity reflects biological/experimental variability rather than poor exponential fits. Three feature relationships were consistent across all panels and both datasets: peak sharpness and peak width were negatively correlated, peak width and inflection time were positively correlated, and peak sharpness and inflection time were negatively correlated, such that spikes with earlier inflection points tend to have wider, less sharp peaks. In pvc-6 (**a-b**), additional strong correlations involving the exponential decay rate (λ) and peak features were present. These pvc-6 patterns were recapitulated in individual spe-1 cells (**c**), but when averaged across the spe-1 population (**d**), most correlations attenuated to near zero, with only the peak sharpness/width/inflection time cluster remaining significant. Panel **e** shows that this attenuation reflects genuine heterogeneity in exp λ correlation direction across cells, rather than poor waveform fitting.

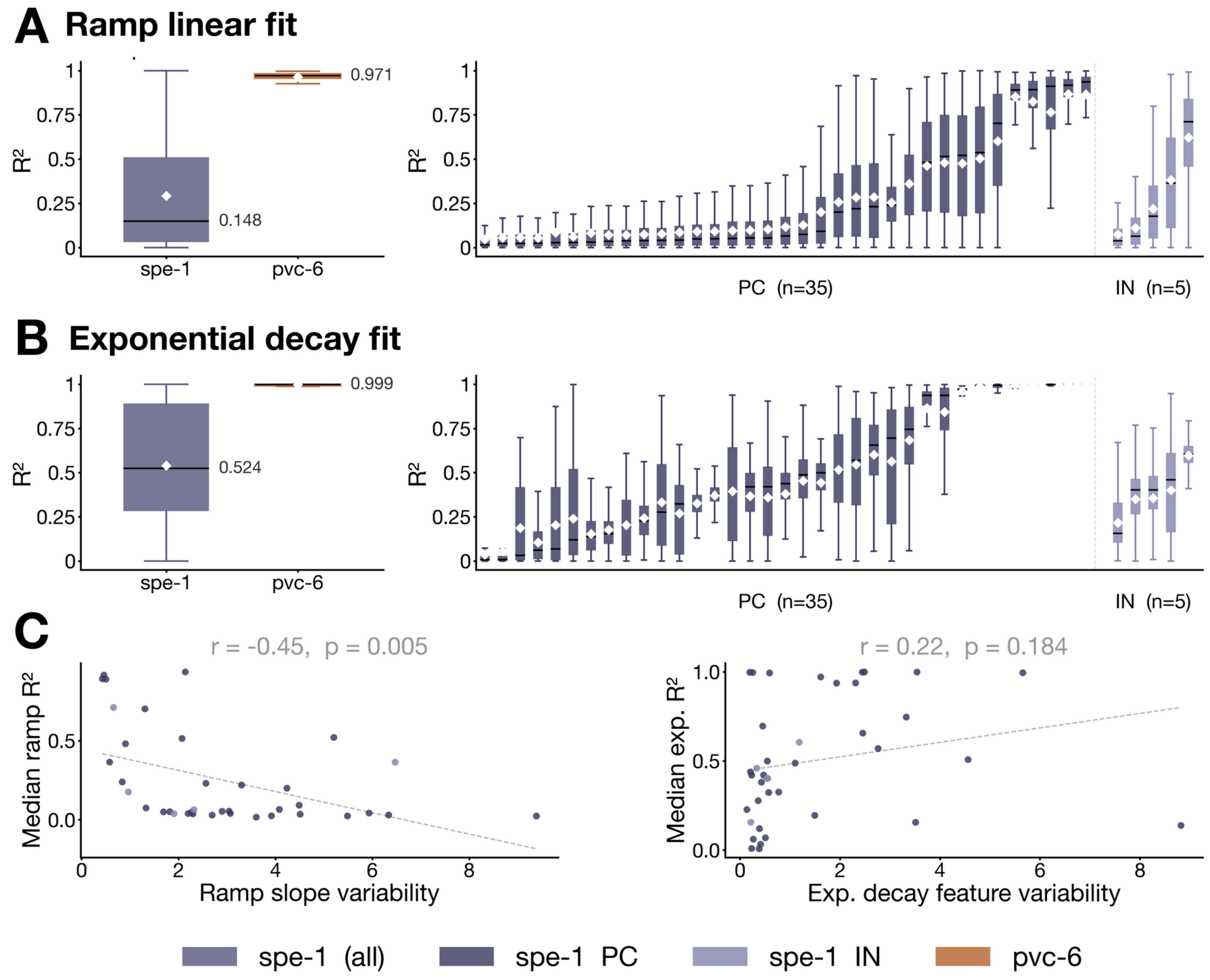


**Supp. Fig. 2 | Spike waveform ramp and decay fits.** Left: Distribution of ramp linear fit $R^2$ across all spikes pooled within spe-1 (purple, median = 0.148) and pvc-6 (brown, median = 0.971). Boxes show interquartile range; whiskers extend to 1.5×IQR; white diamonds indicate means. Right: Per-cell ramp linear fit $R^2$ for spe-1 putative pyramidal cells (PC, dark purple, n=35) and putative interneurons (IN, light purple, n=5), sorted by median $R^2$. **b**, Same as **a** for exponential decay fit $R^2$ (spe-1 median = 0.524; pvc-6 median = 0.999). **c**, (C) Per-cell relationship between fit quality (median $R^2$) and variability of the features extracted from each fit (σ / |median|)| across spikes within each cell): ramp amplitude for the ramp linear fit (left), and exp_lambda and exp_const for the exponential decay fit (right). For the exponential decay, no significant relationship was observed (r = 0.22, p = 0.184), indicating that variability in decay features is independent of fit quality. For the ramp, cells with more variable ramp amplitudes tended to have lower ramp $R^2$ (r = −0.45, p = 0.005). This likely reflects that greater genuine variability in pre-spike ramp dynamics across spikes makes those spikes harder to capture with a single linear model, rather than indicating that poor fits are artificially inflating feature variability. Cells where σ / |median| exceeded 10 were excluded (ramp: n=1; exp: n=2) for visualization.

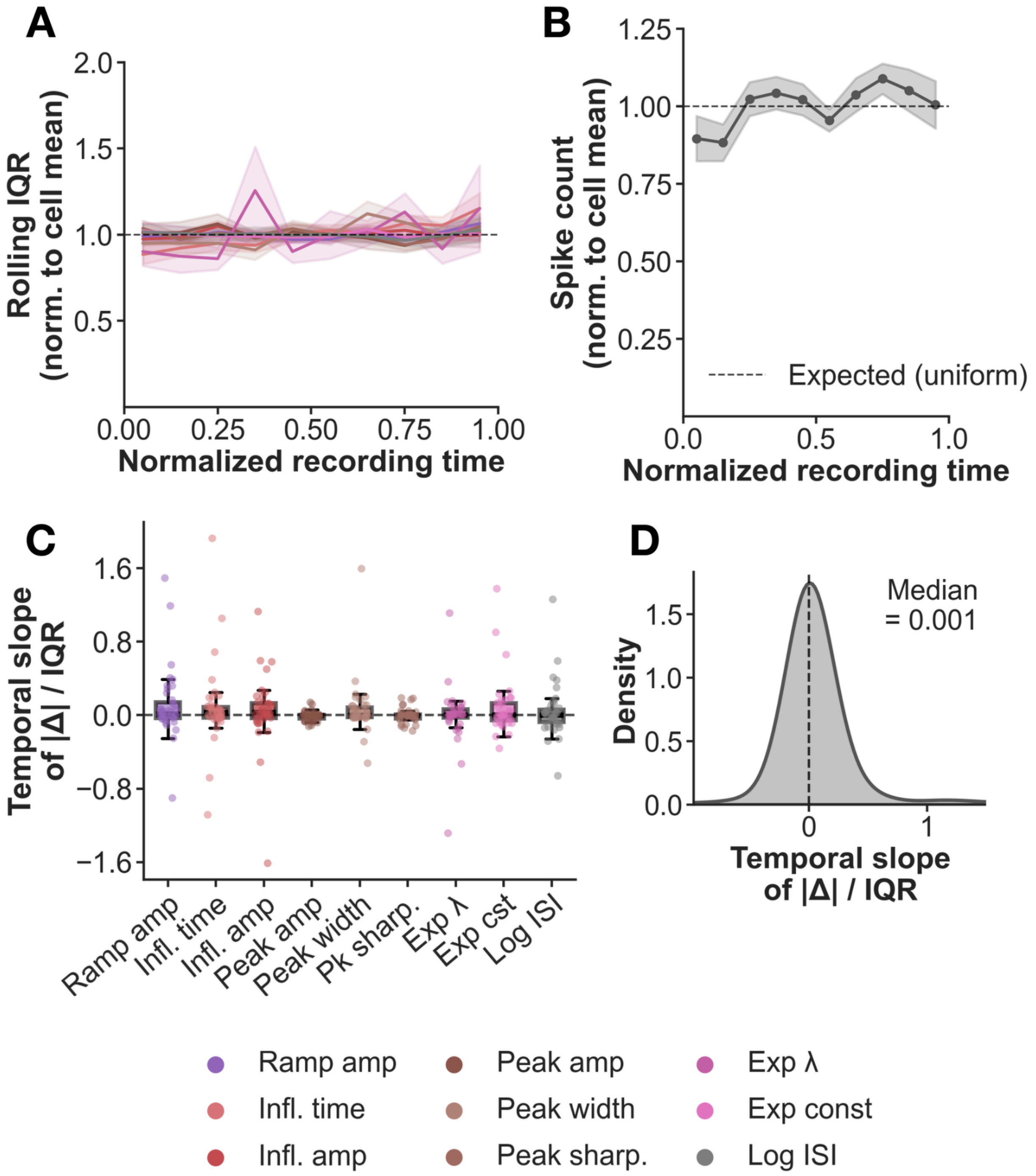


**Supp. Fig. 3 | AP waveform parameter variability is stable across recording time. a**, Rolling IQR of each AP waveform feature normalized to each cell's own mean IQR, averaged across cells (mean ± SEM). A flat profile at 1.0 indicates that within-cell parameter variability is constant throughout the recording, with no evidence of increasing spread over time. **b**, Spike count per normalized time bin, normalized to each cell's mean count (mean ± SEM). The flat profile near the expected uniform rate (dashed line) indicates cells maintained stable firing throughout recordings. **c**, For each cell × feature pair, we fit a linear regression of absolute deviation from the cell mean (normalized by population IQR) against normalized recording time. A positive slope would indicate growing parameter variability (signature of patch degradation). Slopes are symmetrically distributed around zero across all features. **d**, Distribution of temporal slopes pooled across all cell × feature pairs (median = 0.001). The distribution is sharply peaked at zero with no positive skew, indicating no systematic increase in waveform parameter variability over the course of recordings.

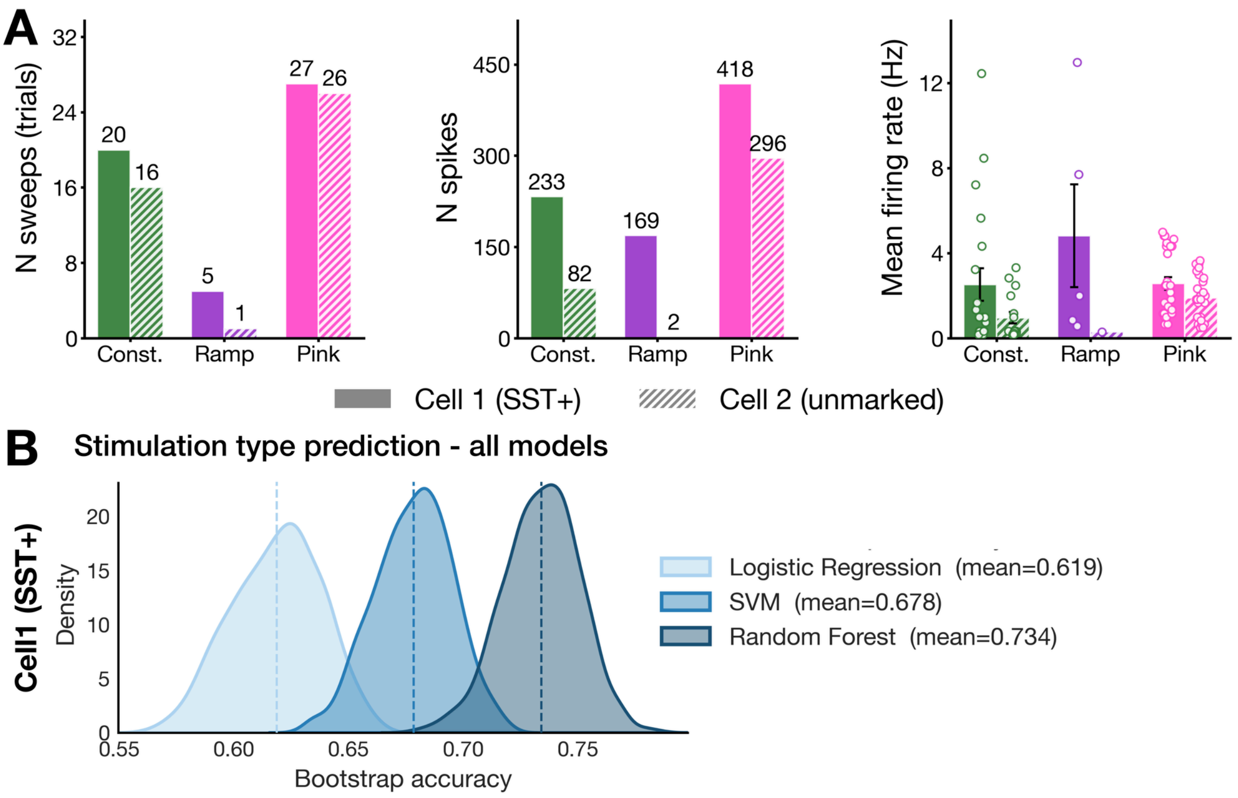


**Supp. Fig. 4 | pvc-6 patch stimulation dataset: full model comparison for stimulation type prediction. a**, Recording statistics for both pvc-6 (Berg, 2014) neurons across three current injection types (constant, ramp, and pink noise). Bar charts show the number of sweeps (trials; left), total spike count (middle), and mean firing rate in Hz (right) for Cell 1 (SST+, solid bars) and Cell 2 (unmarked, hatched bars). In the firing rate panel, individual trial values are overlaid as circles; error bars indicate mean ± SD. Colors denote stimulation type: green = constant, purple = ramp, magenta = pink noise. **b**, Bootstrap accuracy distributions for three-class stimulation type prediction (constant vs. ramp vs. pink noise) from spike waveform features for Cell 1 (SST+) only. Each distribution reflects 1000 bootstrap iterations for logistic regression (light blue; mean = 0.619), support vector machine (SVM; medium blue; mean = 0.678), and random forest (dark blue; mean = 0.734). Dashed vertical lines indicate the mean accuracy for each model. Chance level is 0.33 (three-class classification). Stimulation type prediction was not performed for Cell 2 due to insufficient ramp trials (1 sweep, 2 spikes; see **a**) and class imbalance across conditions (82 constant vs. 296 pink noise spikes), which together precluded reliable multi-class classification.

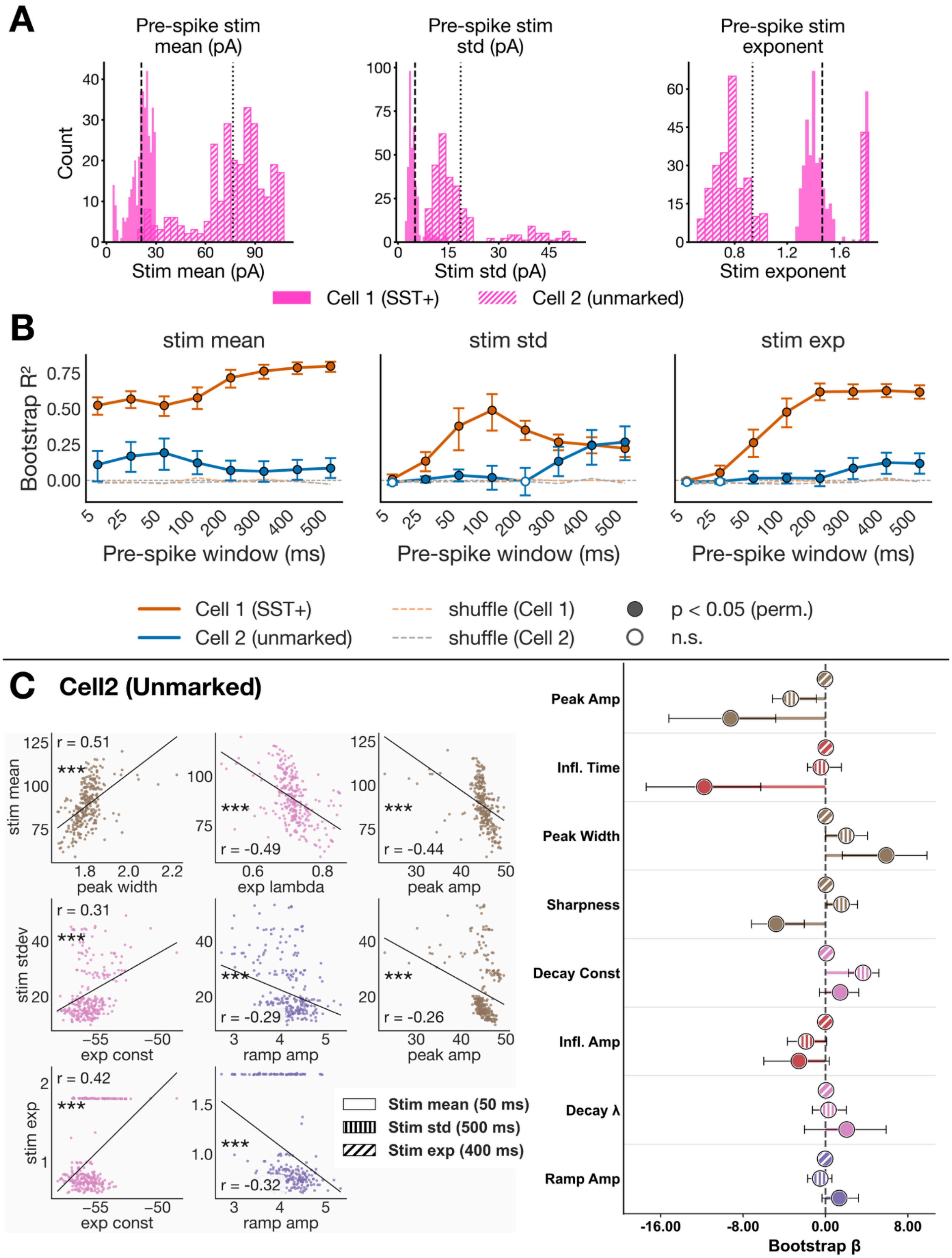
A
Pre-spike stim mean (pA)
Count
Stim mean (pA)
Pre-spike stim std (pA)
Stim std (pA)
Pre-spike stim exponent
Stim exponent
Cell 1 (SST+)
Cell 2 (unmarked)
B
stim mean
stim std
stim exp
Bootstrap R²
Pre-spike window (ms)
Cell 1 (SST+)
Cell 2 (unmarked)
shuffle (Cell 1)
shuffle (Cell 2)
p < 0.05 (perm.)
n.s.
C
Cell2 (Unmarked)
stim mean
stim stdev
stim exp
peak width
exp lambda
peak amp
exp const
ramp amp
r = 0.51
r = -0.49
r = -0.44
r = 0.31
r = -0.29
r = -0.26
r = 0.42
r = -0.32
Stim mean (50 ms)
Stim std (500 ms)
Stim exp (400 ms)
Peak Amp
Infl. Time
Peak Width
Sharpness
Decay Const
Infl. Amp
Decay λ
Ramp Amp
Bootstrap β

**Supp. Fig. 5 | pvc-6 patch stimulation dataset: ridge regression for pink noise stimulus statistics prediction. a**, Distributions of pink noise stimulus statistics across all spikes for Cell 1 (SST+, solid bars) and Cell 2 (unmarked, hatched bars), computed in a 200 ms pre-spike window. Panels show stimulus mean (pA), stimulus standard deviation (pA), and spectral exponent. Dashed vertical lines indicate the per-cell mean (Cell 1, dashed; Cell 2, dotted). **b**, Ridge regression $R^2$ as a function of pre-spike stimulus window length (5–500 ms) for Cell 1 (SST+) and Cell 2 (unmarked). For each window, spike waveform features predicted each stimulus feature via ridge regression with k-fold cross-validation; points show bootstrap mean $R^2 \pm 95\%$ CI. Filled circles indicate windows significantly exceeding a permutation null ($p < 0.05$); open circles indicate non-significant windows. Dashed lines show shuffle controls in which the correspondence between spike waveform features and stimulus statistics was randomly broken by shuffling stimulus labels across spikes (1000 permutations; Cell 1, gray; Cell 2, light gray). Cell 1 (SST+) shows substantially higher $R^2$ than Cell 2 (unmarked) across all stimulus statistics and window lengths. **c**, Spike waveform prediction of stimulus feature for Cell 2 (unmarked) at per-feature optimal windows. Left: Pearson correlation scatter plots at the independently selected best pre-spike window for each stimulus feature (stim mean: 50 ms; stim std: 500 ms; stim exp: 400 ms). Only statistically significant pairs are shown, ranked by $|r|$; Pearson r and significance (FDR corrected, *** $p < 0.001$) are annotated in each panel. Right: Ridge regression beta weights for significant waveform feature–stimulus statistic pairs at their respective best windows (one-sample t-test against zero, FDR corrected ***$p < 0.001$); fill pattern indicates stimulus target (plain = stim mean; vertical lines = stim std; diagonal lines = stim exp). Error bars indicate 95% bootstrap CIs; dashed vertical line at zero.

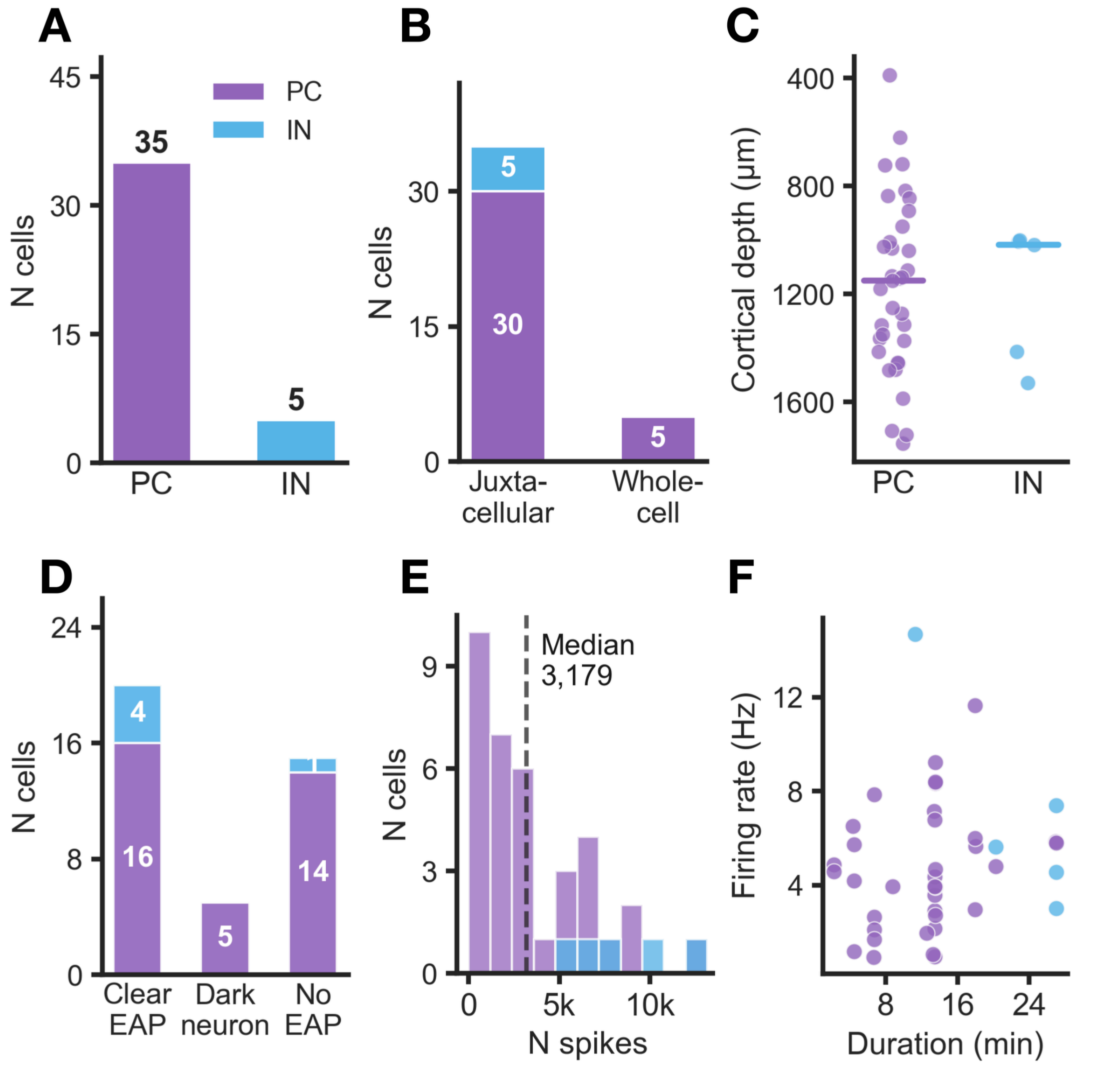


**Supp. Fig. 6 | spe-1 simultaneous patch and neuropixel dataset: Overview of neurons in dataset.** Summary of the n = 40 neurons included in the waveform clustering and LFP analyses, from the spe-1 dataset (Marques-Smith et al., 2020): simultaneous juxtacellular or whole-cell patch-clamp and Neuropixels LFP recordings in rat primary motor and somatosensory cortex under urethane anaesthesia. **a**, Number of cells by putative cell type. Pyramidal cells (PC) and interneurons (IN) were classified from the extracellular spike waveform revealed by the patch spike-triggered average (PSTA) of the Neuropixels channel with the highest peak-peak amplitude, using trough-to-peak duration and negative peak half-width as features. **b**, Number of cells by recording modality (juxtacellular cell-attached vs. whole-cell patch-clamp). **c**, Cortical depth of each cell, estimated from the Bonsai-tracked probe manipulator position (depth of the Neuropixels channel closest to the patch pipette tip). Horizontal bars show the median per cell type; depth axis is inverted. **d**, EAP detectability on the Neuropixels probe, assessed via the PSTA of the extracellular voltage (4-ms windows centred on each patch-clamp spike). “Clear EAP”: canonical waveform with peak-peak amplitude >10 μV; “Dark neuron”: no detectable PSTA waveform despite proximity to the probe and session conditions expected to yield detection; “No EAP”: absence attributed to technical factors (probe drift or misalignment). **e**, Distribution of spike counts per cell. Dashed line shows the median (3,179 spikes). **f**, Recording duration vs. mean firing rate per cell. Colors: purple = PC, blue = IN.

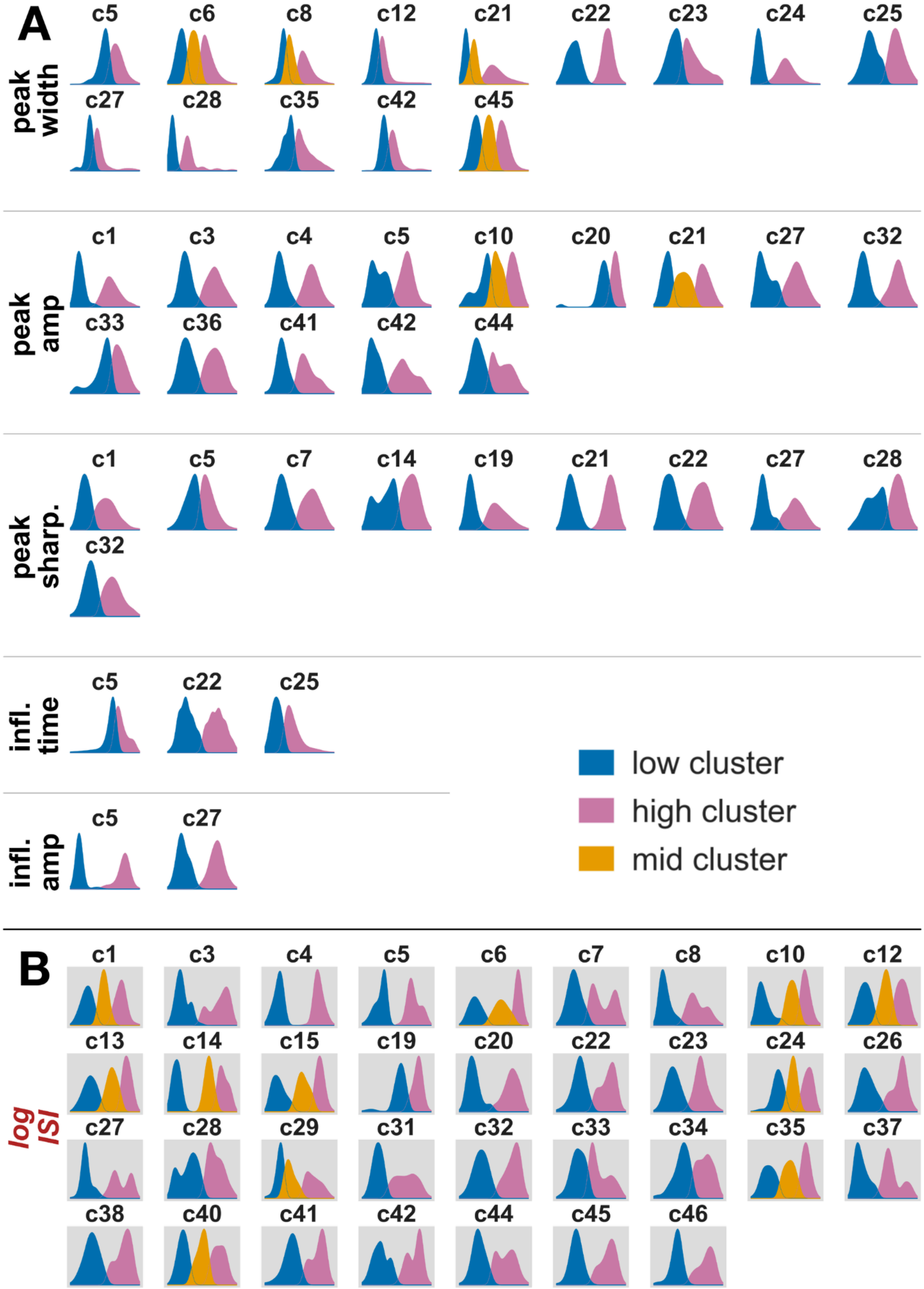
A
peak width
c5 c6 c8 c12 c21 c22 c23 c24 c25
c27 c28 c35 c42 c45
peak amp
c1 c3 c4 c5 c10 c20 c21 c27 c32
c33 c36 c41 c42 c44
peak sharp.
c1 c5 c7 c14 c19 c21 c22 c27 c28
c32
infl. time
c5 c22 c25
infl. amp
c5 c27
low cluster
high cluster
mid cluster
B
log ISI
c1 c3 c4 c5 c6 c7 c8 c10 c12
c13 c14 c15 c19 c20 c22 c23 c24 c26
c27 c28 c29 c31 c32 c33 c34 c35 c37
c38 c40 c41 c42 c44 c45 c46

**Supp. Fig. 7 | spe-1 simultaneous patch and neuropixel dataset: within-neuron spike feature distributions. a**, Within-cell distributions of spike waveform features for all spe-1 neurons in which ≥2 clusters were detected. Clusters represent groups of spikes with distinct feature values within a single neuron, identified via histogram peak analysis (see Methods). Each panel shows one neuron's spike distribution split by cluster assignment. Colors indicate cluster rank: blue = low, pink = high, gold = mid (three-cluster cells only). **b**, Same as A for log inter-spike interval (log ISI; grey background)

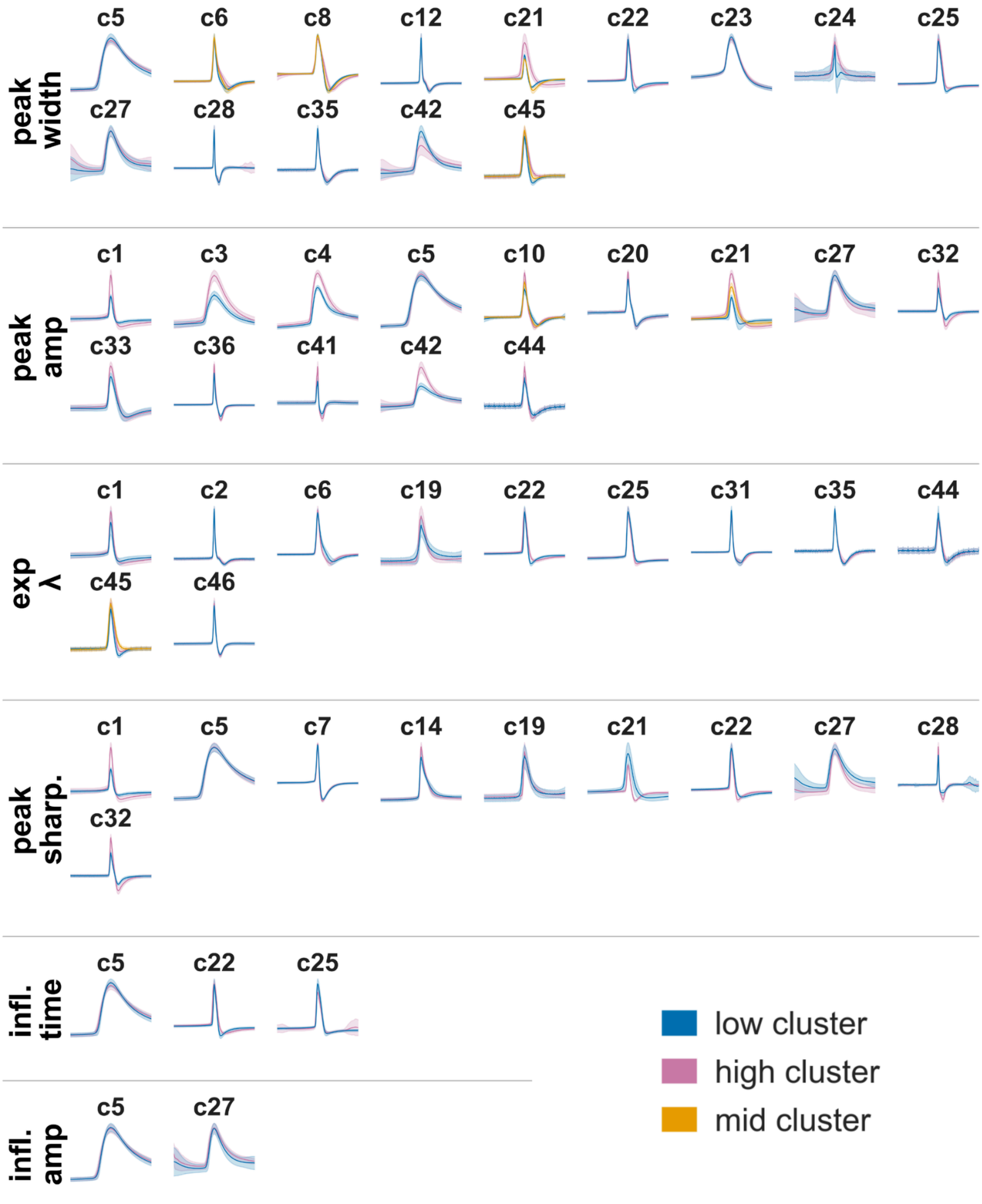


**Supp. Fig. 8 | spe-1 simultaneous patch and neuropixel dataset: within-neuron mean spike waveforms by feature cluster.**

Each panel shows the mean spike waveforms (± 1 SD, shaded) per cluster for one neuron, aligned to the waveform peak. Rows group neurons by which feature was clustered. Only neurons with ≥2 detected clusters for a given feature are shown. Colors indicate cluster rank : blue = low, pink = high, gold = mid (three-cluster cells only). Cluster assignments match those in **Supp. Fig. 7**.

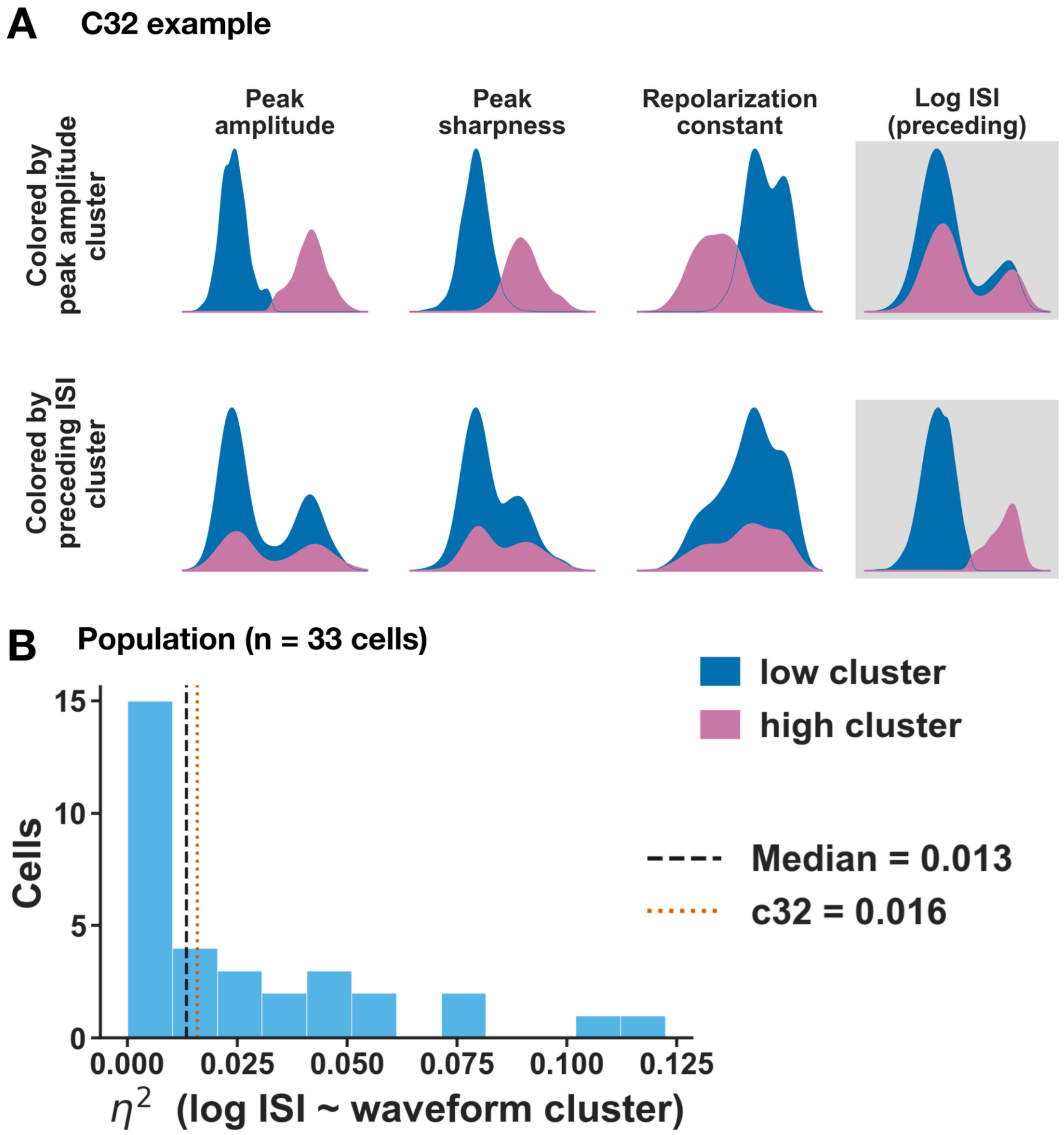


**Supp. Fig. 9 | Spike waveform clusters are independent of ISI-based clusters. a**, Feature distributions for an example cell (c32, n = 2887 spikes), shown as solid-fill kernel density estimates (IQR × 3 outlier removal, Scott's rule bandwidth, proportional normalization). Row 1: spikes colored by peak amplitude cluster assignment (low cluster, blue; high cluster, pink). Waveform features (peak amplitude, peak sharpness, repolarization constant) are clearly separated between cluster groups; log inter-spike interval (log ISI; grey background) is not. Row 2: same spikes colored by ISI cluster assignment. Log ISI is clearly separated; waveform features are not. **b**, Population distribution of $\eta^2$ (Kruskal-Wallis H / (n − 1)) quantifying the association between waveform cluster assignment and log ISI across all cells with at least one waveform cluster and an ISI cluster (n = 33 cells). Dashed line: population median. Dotted line: c32. $\eta^2$ near 0 indicates that waveform cluster membership does not explain ISI variability.

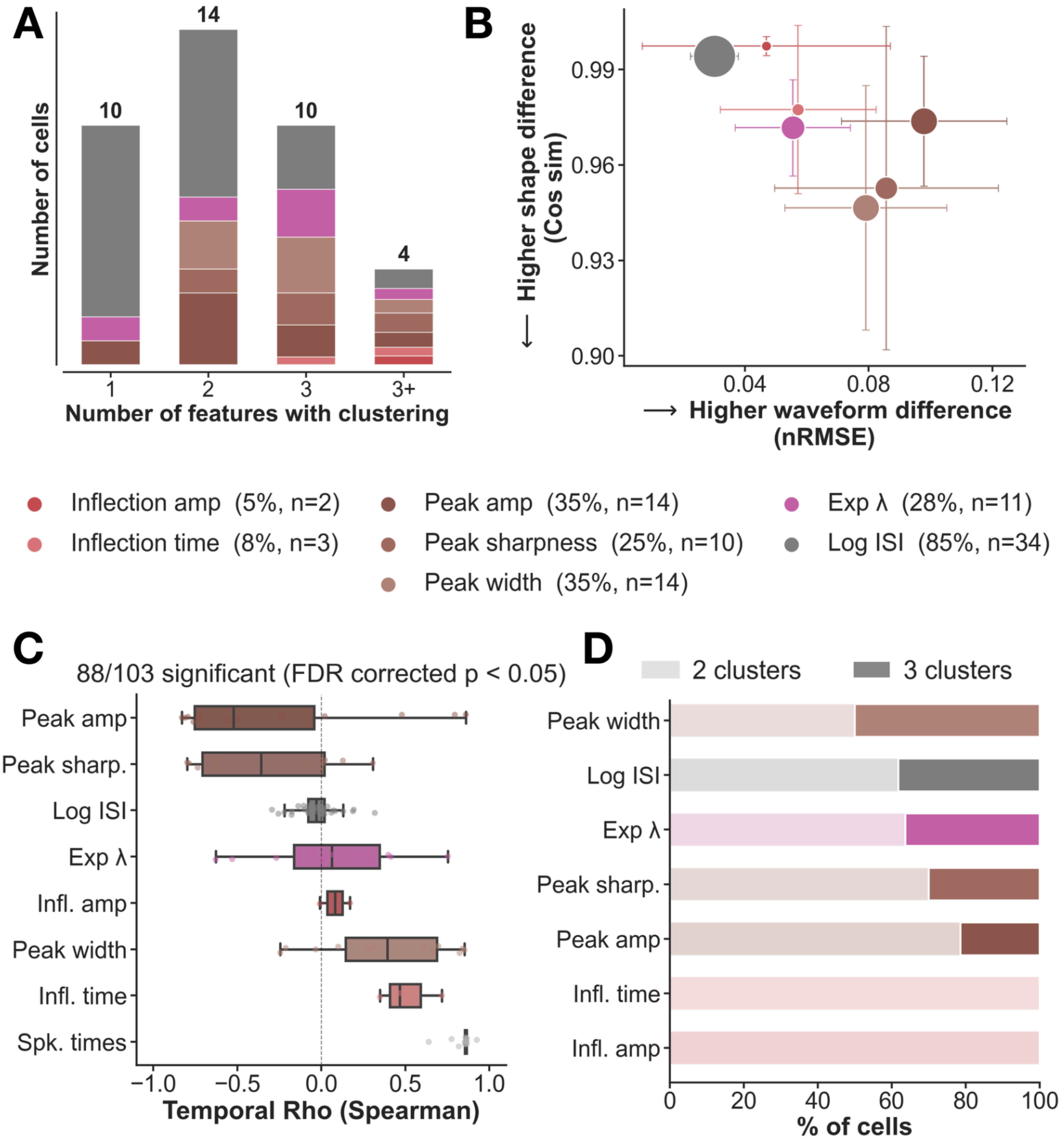


**Supp. Fig. 10 | spe-1 simultaneous patch and neuropixel dataset: Characterization of spike waveform cluster properties across cells. a**, Number of cells exhibiting multimodal clustering in 1, 2, 3, or 3+ waveform features. Bar segments are colored by feature (see legend). **b**, Mean nRMSE (x-axis;

normalized root mean squared error, capturing overall waveform difference including amplitude) versus cosine similarity (y-axis; a measure of waveform shape similarity independent of amplitude) between waveform clusters per feature. Marker size is proportional to the percentage of cells showing clustering for that feature (see legend). Error bars indicate ±1 SD across cells. **c**, Temporal structure of cluster membership. Spearman ρ between spike index and ordinal cluster label quantifies whether waveform cluster identity changes systematically over the course of a recording — i.e., whether waveform shape is stationary or changes through recording time. High |ρ| indicates that spikes recorded at different time points tend to belong to different clusters. 88/103 cell × feature pairs showed significant temporal structure (FDR corrected $p < 0.05$). **d**, Fraction of cells with 2 (light) versus 3 (solid) total waveform clusters per spike feature. Features are sorted by ascending 3-cluster fraction.

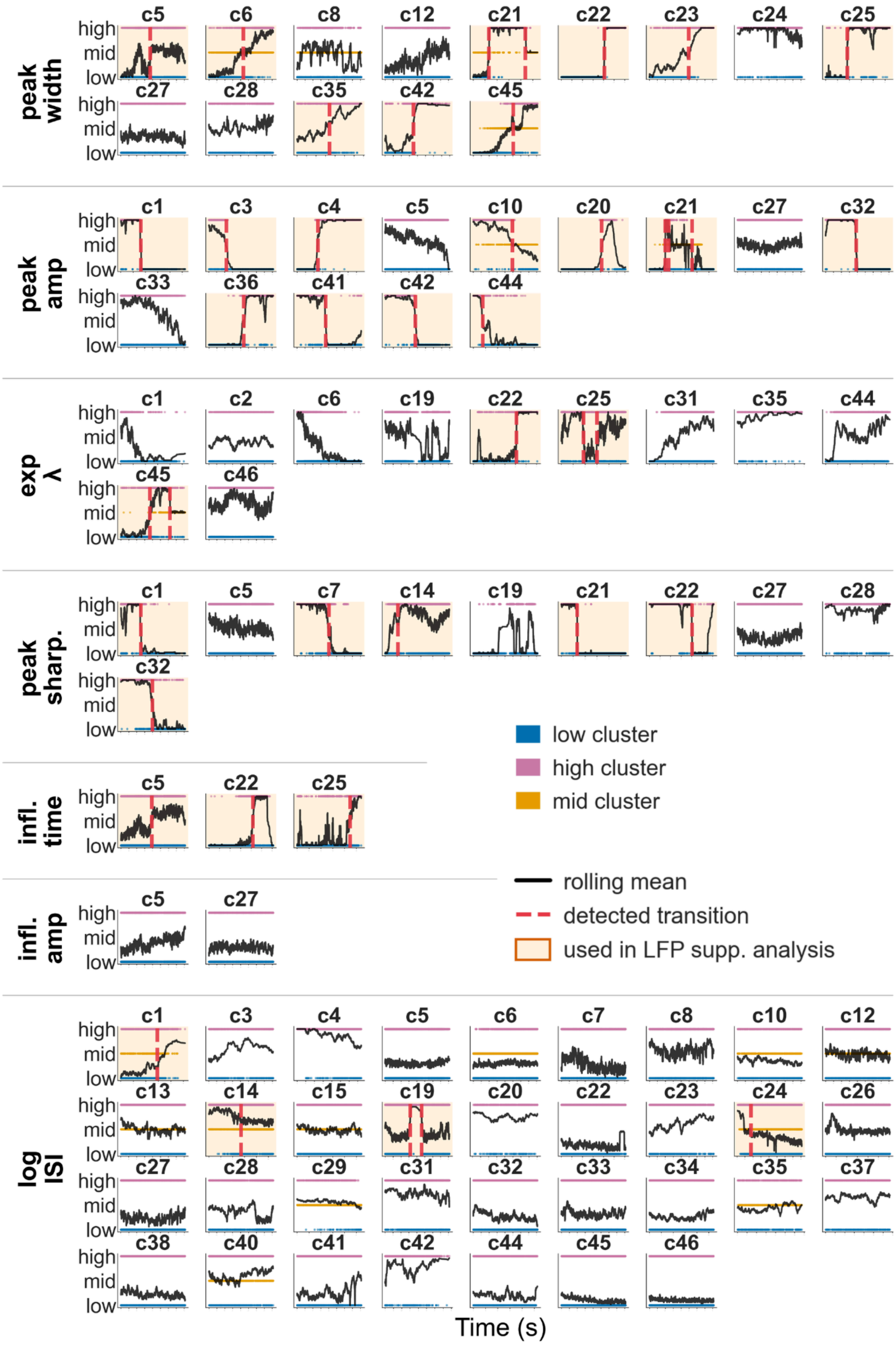

peak width
c5 c6 c8 c12 c21 c22 c23 c24 c25
c27 c28 c35 c42 c45
peak amp
c1 c3 c4 c5 c10 c20 c21 c27 c32
c33 c36 c41 c42 c44
exp λ
c1 c2 c6 c19 c22 c25 c31 c35 c44
c45 c46
peak sharp.
c1 c5 c7 c14 c19 c21 c22 c27 c28
c32
infl. time
c5 c22 c25
infl. amp
c5 c27
log ISI
c1 c3 c4 c5 c6 c7 c8 c10 c12
c13 c14 c15 c19 c20 c22 c23 c24 c26
c27 c28 c29 c31 c32 c33 c34 c35 c37
c38 c40 c41 c42 c44 c45 c46
high
mid
low
low cluster
high cluster
mid cluster
rolling mean
detected transition
used in LFP supp. analysis
Time (s)

**Supp. Fig. 11 | spe-1 simultaneous patch and neuropixel dataset: temporal trajectories of within-neuron spike feature.** Each panel shows how a single cell's cluster membership for one spike feature evolves over the course of the recording. Colored dots represent individual spikes color-coded by cluster assignment (blue = low, pink = high, gold = mid); the black line shows the rolling mean of ordinal cluster labels (window = 50 spikes), and the x-axis spans the full recording duration. Vertical dashed red lines indicate detected transition times, identified by fitting a multi-component sigmoid model to the rolling mean. Panels highlighted in amber are cell × feature pairs included in the peri-transition LFP spectral analysis (**Supp. Fig. 12**), in which LFP windows were extracted around each detected transition.

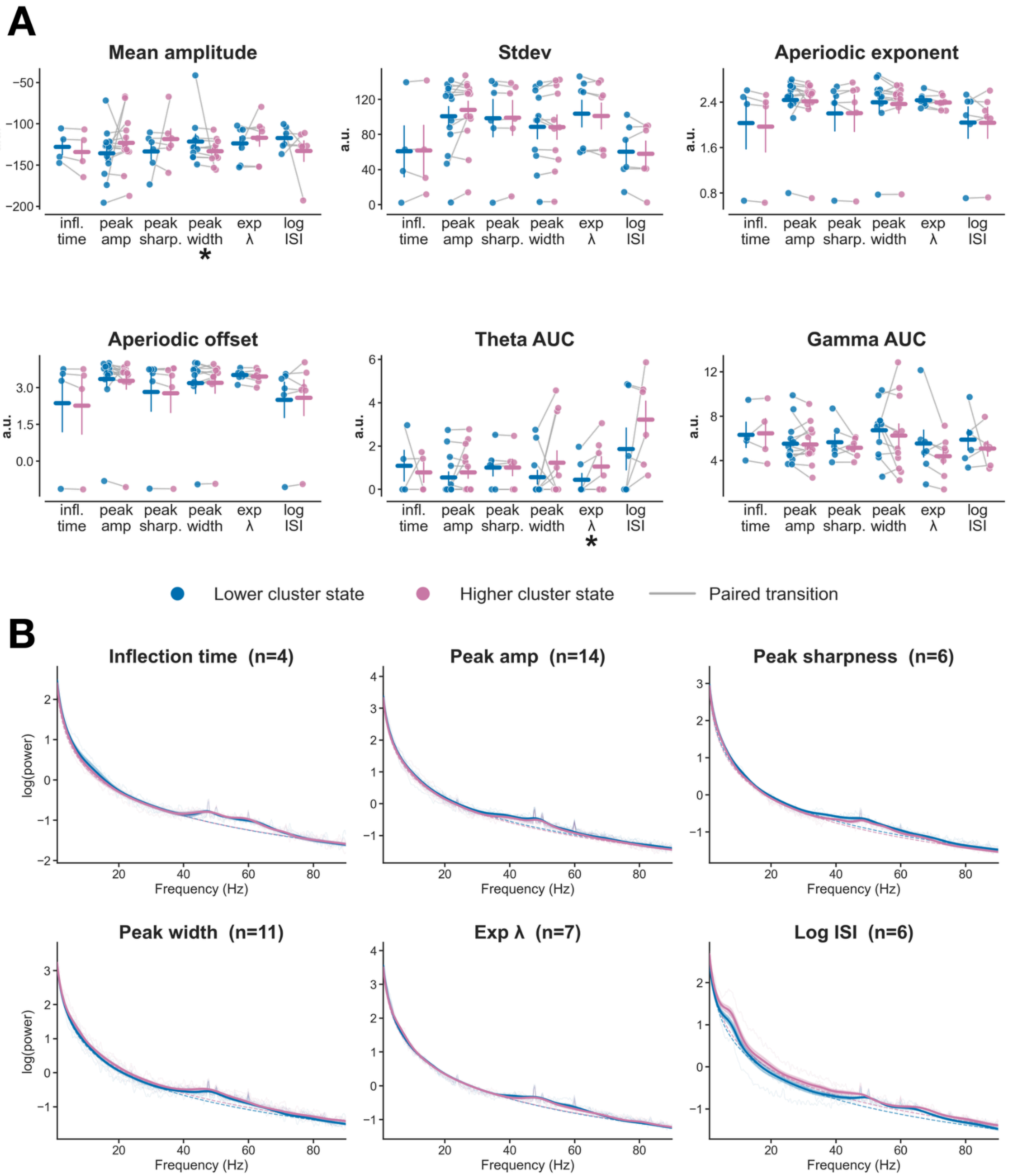


**Supp. Fig. 12 | spe-1 simultaneous patch and neuropixel dataset: LFP dynamics during spike waveform cluster transitions. a**, LFP scalar metrics comparing lower vs. higher spike cluster states across spike waveform features. Transitions were identified as described in **Supp. Fig. 11** (sigmoid fits to within-neuron spike feature trajectories; $R^2 \geq 0.80$). To pool across transition directions, LFP blocks were sign-corrected such that the lower cluster state always corresponds to the pre-transition block for low→high transitions and the post-transition block for high→low transitions (and vice versa for the higher cluster state). Each panel shows one LFP metric; within each panel, paired dots (blue = lower cluster

state, pink = higher cluster state) are plotted for each spike waveform feature (x-axis). Connected gray lines link paired measurements from the same transition event. Horizontal bars and vertical lines denote mean ± SEM. Mean LFP amplitude was significantly lower during the higher cluster state for peak width-defined transitions. Theta AUC was significantly higher during the higher cluster state for exp λ-defined transitions. (Wilcoxon signed-rank test; * $p < 0.05$, ** $p < 0.01$, *** $p < 0.001$). Aperiodic exponent ($\mu V^2\ Hz^{-1}$) and offset ($\mu V^2$) are derived from specparam fits to the LFP power spectrum (fitting range: 1–90 Hz). Theta AUC and Gamma AUC represent the area under the periodic component of the power spectrum in the theta (4–15 Hz) and gamma (30–80 Hz) bands, respectively (arbitrary units; see Methods). **b**, Mean LFP power spectra for lower (blue) and higher (pink) cluster states, shown separately for each spike waveform feature, using the same sign-correction as in (A). Thin lines show individual-transition raw power spectral densities; dashed lines show the mean aperiodic fit; solid lines with shading show the mean specparam model fit ± SEM across transitions. n = number of transitions per feature. Panel titles indicate significant band-specific differences where present (Wilcoxon signed-rank test, $p < 0.05$).
Given the small per-feature sample sizes and high inter-transition variability, these results are intended as exploratory rather than definitive evidence of LFP state changes.

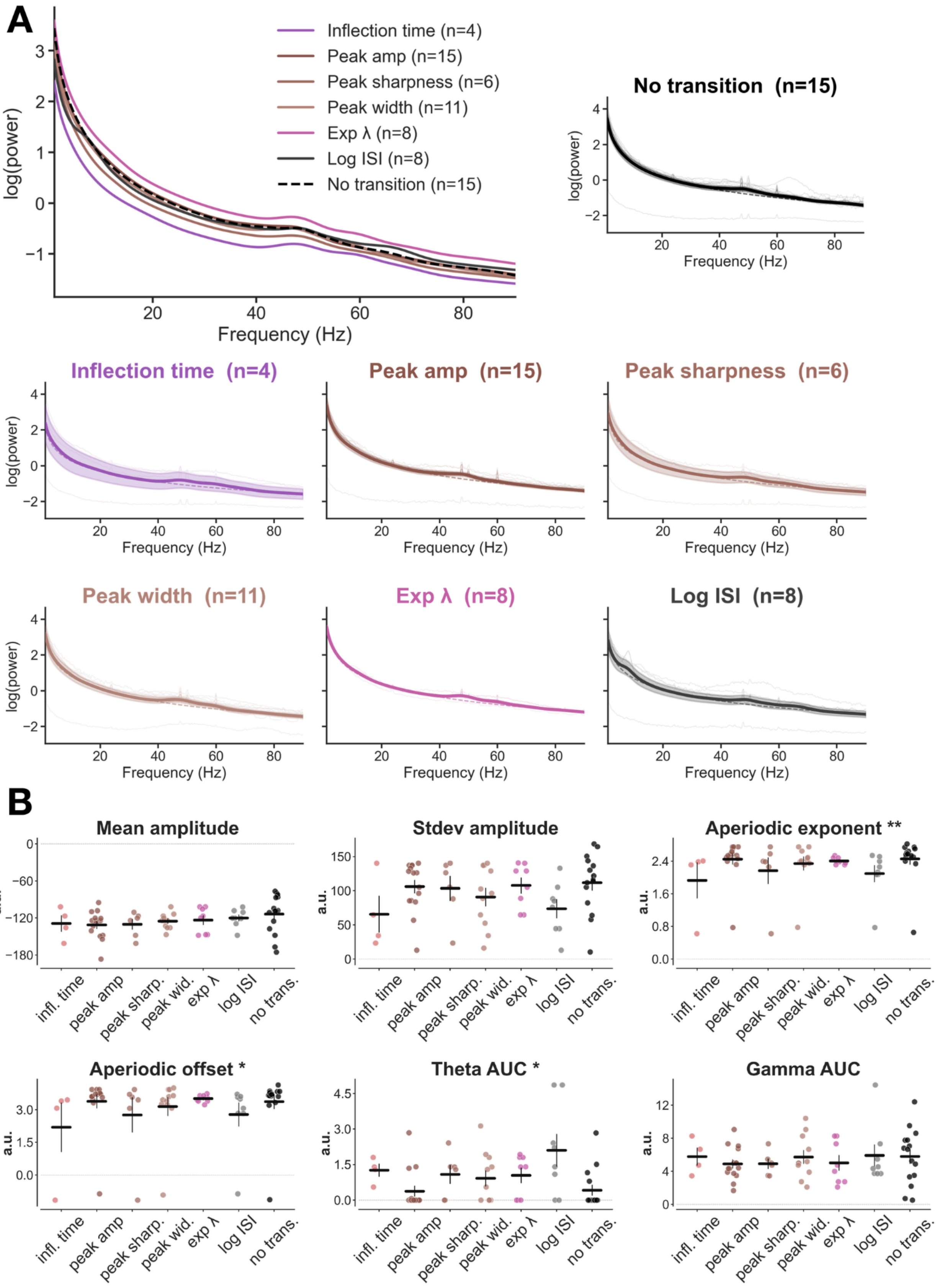

A
Inflection time (n=4)
Peak amp (n=15)
Peak sharpness (n=6)
Peak width (n=11)
Exp λ (n=8)
Log ISI (n=8)
No transition (n=15)
log(power)
Frequency (Hz)
No transition (n=15)
Inflection time (n=4)
Peak amp (n=15)
Peak sharpness (n=6)
Peak width (n=11)
Exp λ (n=8)
Log ISI (n=8)
B
Mean amplitude
Stdev amplitude
Aperiodic exponent **
Aperiodic offset *
Theta AUC *
Gamma AUC
a.u.
infl. time
peak amp
peak sharp.
peak wid.
exp λ
log ISI
no trans.

**Supp. Fig. 13 | spe-1 simultaneous patch and neuropixel dataset: LFP activity by spike feature cluster transitions. a**, LFP power spectra computed over the full recording for each cell, with cells separated by which spike waveform feature exhibits cluster state transitions. Top left: overlay of mean specparam model fits across all feature groups; cells with no detected cluster transitions in any feature are shown as a dashed black reference line. Small panels (top right and bottom grid): per-group mean ± SEM specparam fits (colored lines ± shading), with individual-cell raw power spectral densities shown as thin semi-transparent lines in the background. n = number of cells per group. **b**, LFP metrics for each spike feature group derived from the same whole-recording time domain analysis and specparam fits: mean and standard deviation of broadband amplitude,a periodic exponent and offset, and area under the periodic component in the theta (4–15 Hz) and gamma (30–80 Hz) bands. Each dot represents one cell; crosshairs indicate group mean ± SEM. Stars in panel titles indicate a significant difference across groups (Kruskal–Wallis test; * $p < 0.05$, ** $p < 0.01$). Aperiodic exponent differed significantly across groups ($H(6) = 16.98$, $p = 0.009$), as did aperiodic offset ($H(6) = 13.21$, $p = 0.040$) and theta AUC ($H(6) = 12.73$, $p = 0.048$). Broadband amplitude and gamma AUC did not differ significantly across groups. Given the small per-group sample sizes, these results are intended as exploratory rather than definitive evidence of feature-specific LFP differences, and are not corrected for multiple comparisons across the six metrics tested.

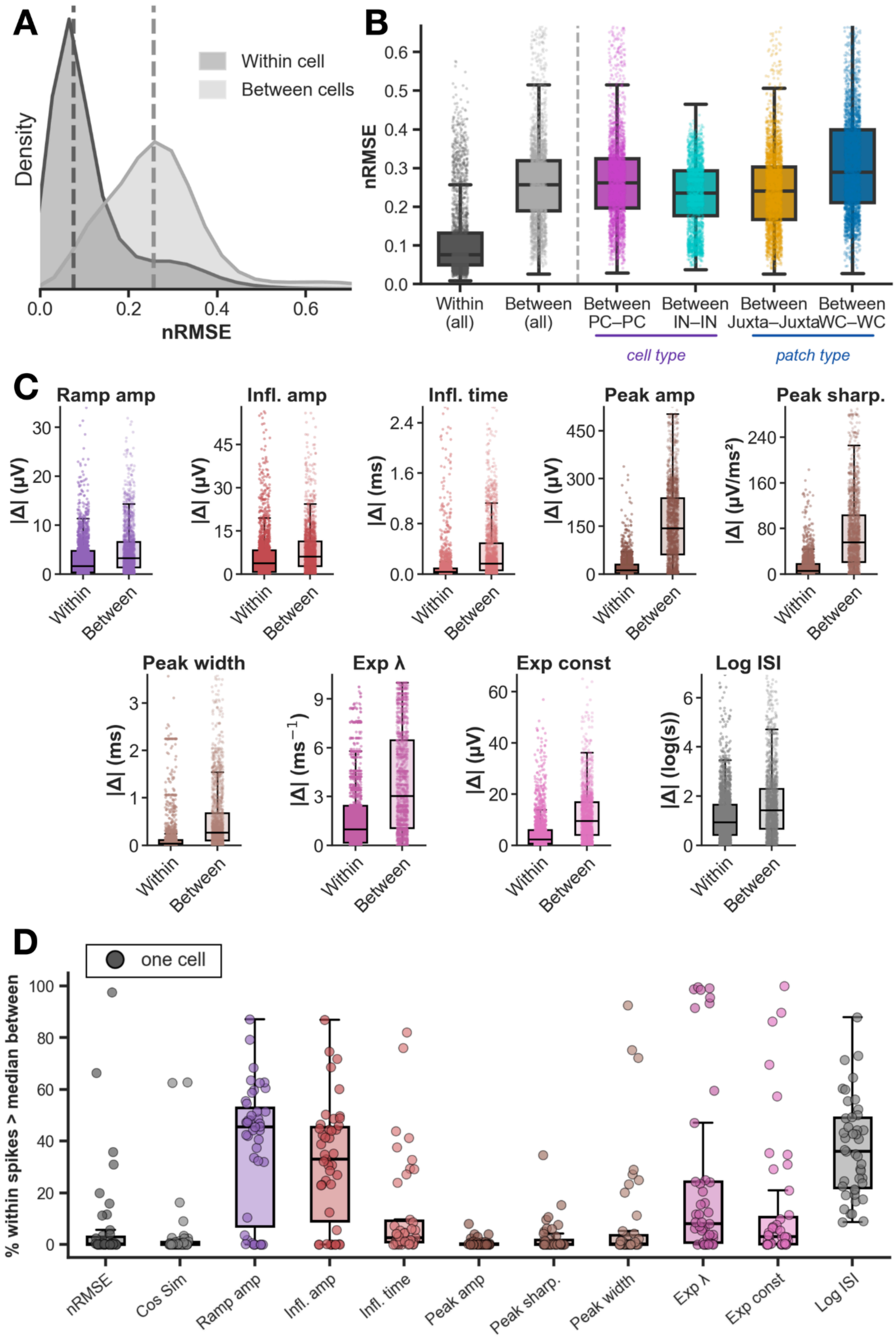
A
Density
nRMSE
Within cell
Between cells
B
nRMSE
Within (all)
Between (all)
Between PC–PC
Between IN–IN
Between Juxta–Juxta
Between WC–WC
cell type
patch type
C
Ramp amp
Infl. amp
Infl. time
Peak amp
Peak sharp.
Peak width
Exp λ
Exp const
Log ISI
|Δ| (µV)
|Δ| (ms)
|Δ| (µV/ms²)
|Δ| (ms⁻¹)
|Δ| (log(s))
Within
Between
D
one cell
% within spikes > median between
nRMSE
Cos Sim
Ramp amp
Infl. amp
Infl. time
Peak amp
Peak sharp.
Peak width
Exp λ
Exp const
Log ISI

**Supp. Fig. 14 | spe-1 simultaneous patch and neuropixel dataset: Within- and between-cell spike waveform variability. a**, Distribution of nRMSE (normalized root mean squared error) comparing each spike to its own cell's mean waveform (within cell, dark) versus to other cells' mean waveforms (between cells, light). nRMSE quantifies waveform amplitude dissimilarity. Dashed lines indicate medians. **b**, nRMSE distributions split by cell-type (PC–PC, IN–IN) and recording-method (Juxta/Juxta, WC/WC) subgroups. Within (all): each spike vs its own cell mean. Between subgroups: spikes compared to mean waveforms of cells within the same subgroup. Dashed vertical line separates within from between comparisons. **c**, Per-feature absolute deviation of individual spike values from their cell's mean (Within) versus from other cells' means (Between), shown in feature units. **d**, Percentage of within-cell spikes whose deviation from the cell mean exceeds the median between-cell distance, per metric and feature (each dot = one cell). Cells near 100% likely reflect strongly bimodal within-cell distributions (i.e., cells with clustering in that feature), where all spikes sit far from the overall cell mean regardless of which cluster they belong to.

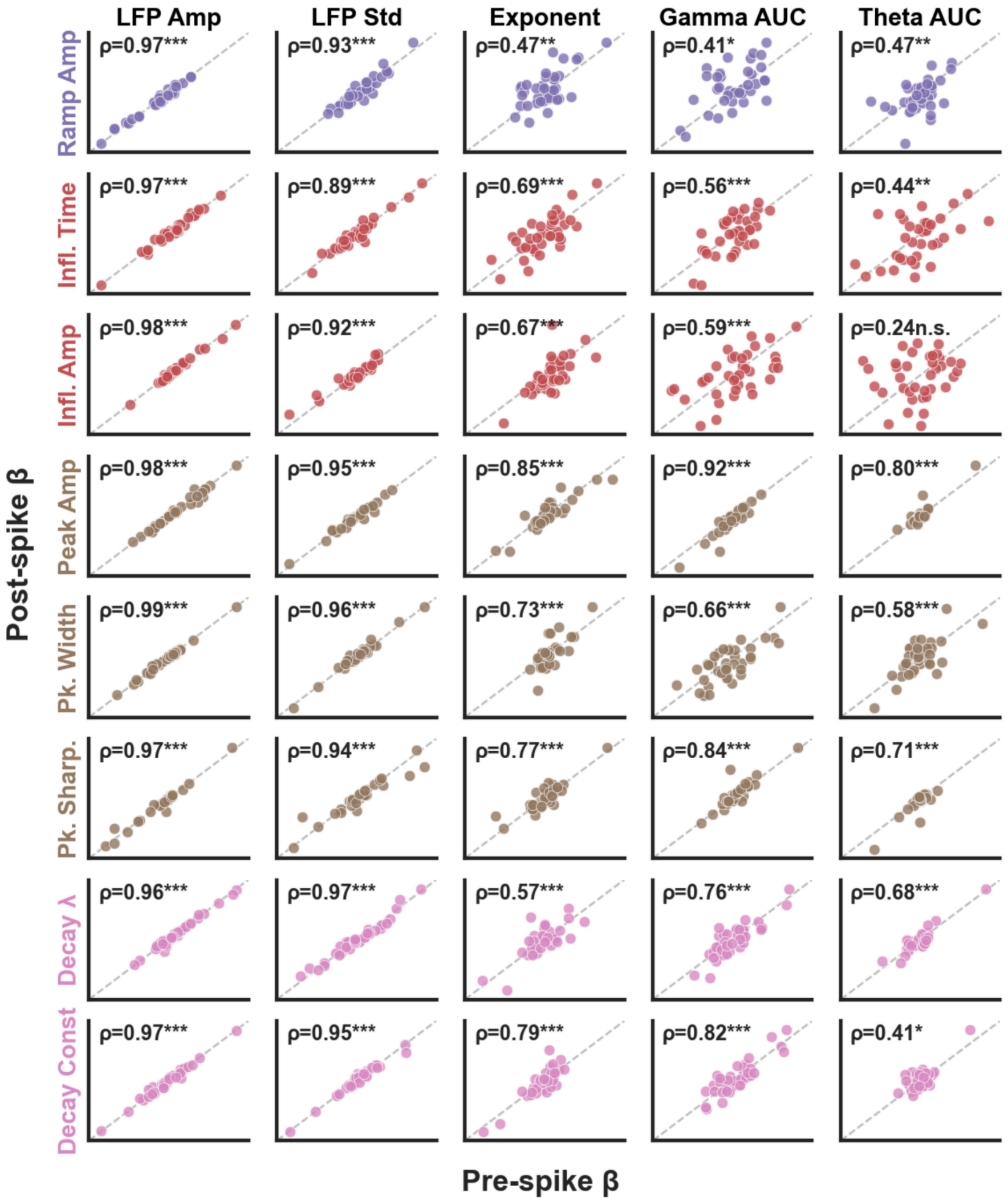


**Supp. Fig. 15 | spe-1 simultaneous patch and neuropixel dataset: Pre- and post-spike regression coefficients are highly correlated across LFP targets and waveform features.** Each panel shows the ridge regression β coefficient estimated from pre-spike LFP windows (x-axis) against the corresponding β from post-spike LFP windows (y-axis), for one waveform feature (rows, color-coded) and one LFP target (columns). Each point is one cell. Dashed line: identity ($y = x$). Spearman ρ and significance are annotated per panel (***$p < 0.001$, **$p < 0.01$, *$p < 0.05$, n.s. $p \geq 0.05$). Coefficients were highly consistent between pre- and post-spike windows for LFP Amp and LFP Std ($\rho \geq 0.89$ for all features), and remained significant for Aperiodic Exponent, Gamma AUC, and Theta AUC, with the exception of Inflection Amplitude × Theta AUC ($\rho = 0.24$, n.s.). Pre- and post-spike $\beta$ coefficients were averaged for all subsequent analyses.

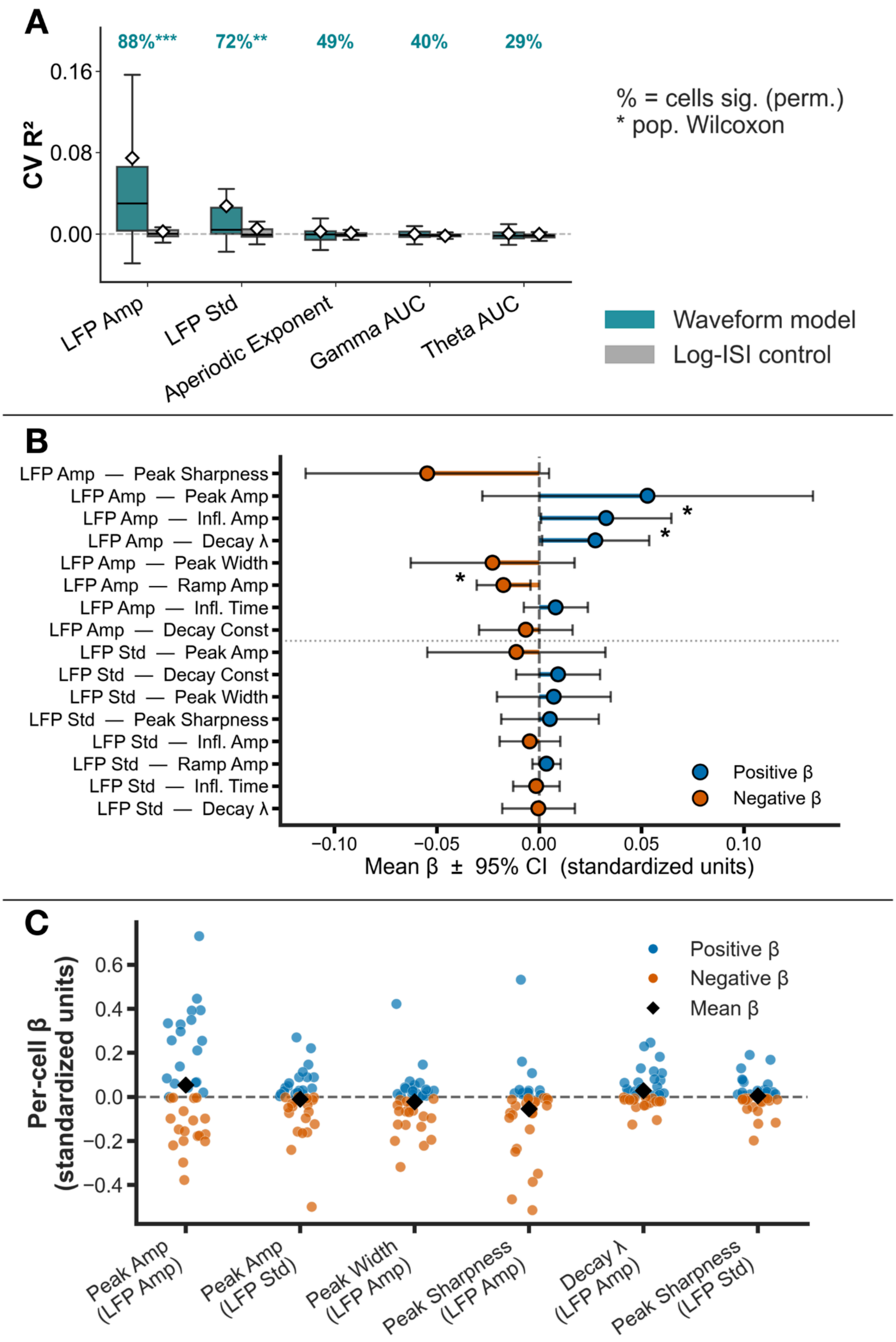
A
88%***
72%**
49%
40%
29%
CV R²
0.16
0.08
0.00
% = cells sig. (perm.)
* pop. Wilcoxon
LFP Amp
LFP Std
Aperiodic Exponent
Gamma AUC
Theta AUC
Waveform model
Log-ISI control
B
LFP Amp — Peak Sharpness
LFP Amp — Peak Amp
LFP Amp — Infl. Amp
LFP Amp — Decay λ
LFP Amp — Peak Width
LFP Amp — Ramp Amp
LFP Amp — Infl. Time
LFP Amp — Decay Const
LFP Std — Peak Amp
LFP Std — Decay Const
LFP Std — Peak Width
LFP Std — Peak Sharpness
LFP Std — Infl. Amp
LFP Std — Ramp Amp
LFP Std — Infl. Time
LFP Std — Decay λ
Positive β
Negative β
−0.10
−0.05
0.00
0.05
0.10
Mean β ± 95% CI (standardized units)
C
Per-cell β (standardized units)
0.6
0.4
0.2
0.0
−0.2
−0.4
Positive β
Negative β
Mean β
Peak Amp (LFP Amp)
Peak Amp (LFP Std)
Peak Width (LFP Amp)
Peak Sharpness (LFP Amp)
Decay λ (LFP Amp)
Peak Sharpness (LFP Std)

**Supp. Fig. 16 | spe-1 simultaneous patch and neuropixel dataset: Waveform shape predicts LFP dynamics: ridge regression model performance and coefficients. a**, Cross-validated $R^2$ (CV $R^2$) for the ridge regression waveform model (teal) and Log-ISI-only control (gray) across five LFP targets, averaged across pre- and post-spike windows. Diamonds indicate the population mean. Percentages above each group indicate the fraction of cells with individually significant models (permutation test, FDR corrected $p < 0.05$); population-level significance among FDR-significant targets (Wilcoxon test, CV $R^2 > 0$, FDR corrected $^{**}p < 0.01$; $^{***}p < 0.001$). **b**, Population-level mean $\beta \pm 95\%$ CI for all 8 waveform features predicting LFP amplitude (top) and LFP standard deviation (bottom; dotted line separates targets). Features sorted by effect size within each target. Asterisks: one-sample t-test, mean $\beta \neq 0$ ($^{*}p < 0.05$; $^{**}p < 0.01$). Pre- and post-spike betas averaged prior to testing (justified by A). Only $R^2$-significant targets shown. **c**, Per-cell $\beta$ distributions for the 6 feature × LFP target combinations with the largest individual cell effects but no population-level agreement, ranked by mean$|\beta|$ × minority-direction fraction. Blue = positive $\beta$, orange = negative $\beta$; diamond = population mean. Parenthetical labels denote LFP target (Amp = LFP amplitude, Std = LFP standard deviation). Despite strong effects in individual cells, these combinations show near-zero population means due to opposing directions across cells.

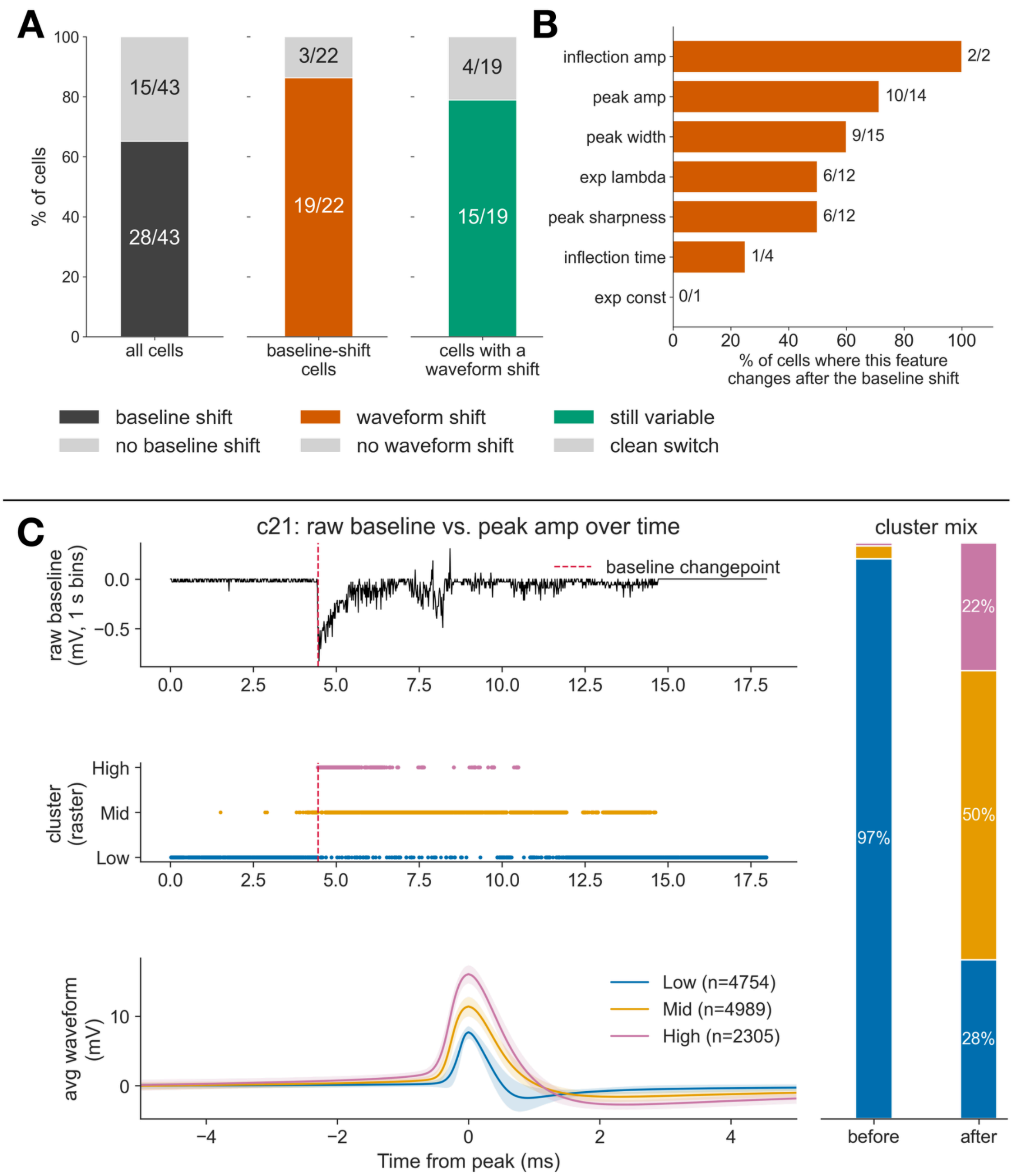


**Supp. Fig. 17 | Baseline membrane-potential shifts frequently coincide with spike waveform shifts.** **a,** Of 43 spe-1 cells, 28 (65%) showed at least one significant shift in baseline membrane potential (PELT changepoint analysis on the raw, unfiltered patch trace; left bar). Of the 22 baseline-shift cells with at least one clustered waveform feature, 19 (86%) also showed a waveform-shape shift, defined as a PELT changepoint in that feature's cluster assignment coinciding with the baseline changepoint (middle bar). Of those 19, 15 (79%) still showed more than one cluster present among spikes following the changepoint rather than a clean switch to a single cluster (right bar). **b**, Fraction of cells, per waveform feature, in which a waveform-cluster changepoint coincided with a baseline changepoint. These baseline shifts may indicate experimenter stimulation, but even so, this does not appear to be the sole driver of the waveform variability reported elsewhere in this manuscript, since most affected cells remained variable rather than switching cleanly to a single new waveform state after the shift. **c**, One example cell (c21) illustrating this pattern: Top: raw baseline trace with the detected changepoint (dashed line). Middle: raster of spikes by peak amplitude cluster, showing the coincident shift. Bottom left: mean spike waveform per cluster. Bottom right: cluster composition before versus after the changepoint. Colors: blue/gold/pink (Low/Mid/High) indicate waveform clusters in **c**.
