## Supplementary material for "Action potential waveforms are state-dependent": Methods and Materials

### MATERIALS AND METHODS

#### OPEN DATA SOURCES

***pvc-6: High-Resolution Controlled Stimulation Dataset***
Whole-cell current clamp recordings from mouse primary visual cortex (V1) were obtained from the CRCNS pvc-6 public dataset (Allen Institute; Berg, 2014). The dataset was downloaded from the Collaborative Research in Computational Neuroscience (CRCNS) database, which additionally describes the data acquisition protocol in detail (Berg, 2014; http://dx.doi.org/10.6080/K0H12ZXD). Whole-cell patch-clamp recordings were obtained from two Layer 2/3 pyramidal neurons in acute coronal brain slices (350 µm) of primary visual cortex from adult mice. Synaptic transmission was blocked with 1 mM kynurenic acid and 0.1 mM picrotoxin. Signals were acquired at 200 kHz and low-pass filtered with a Bessel filter at 10 kHz. Each neuron received three current injection stimulus types: (1) a 500-ms constant-amplitude square pulse at AP threshold, (2) a linearly ramping current, and (3) three variants of 1/f (pink noise), each variant repeated three times. For Cell 1 (SST+ interneuron, rheobase 20 pA), pink noise was presented at 75%, 100%, and 125% of threshold. For Cell 2 (morphologically unmarked, rheobase 60 pA), pink noise was presented at 100%, 125%, and 150% of threshold. Cell 1 is used in all main-text analyses. Cell 2 is presented in supplementary pink-noise analyses only, as it comprised very few constant and ramp sweeps, precluding stimulus-type classification.

***Preprocessing:*** Recordings were loaded directly from CRCNS in their provided HDF5 format. No additional filtering was applied to the patch-clamp signal prior to spike detection, as recordings were already acquired at 200 kHz with a high signal-to-noise ratio. Sweeps were labeled by stimulus type directly from the injected-current trace: the Pearson correlation between current amplitude and sample index was computed after discarding the first 5000 samples; sweeps with ≤2 nonzero samples were classified as "none," those with $r > 0.95$ as "ramp," those with an undefined (NaN) correlation — indicating a flat, constant trace — as "constant," and all others as "pink noise." Spikes were detected as threshold crossings in the voltage trace (threshold: −10 mV) using a 5 ms pre- and 5 ms post-peak extraction window. The last spike of each sweep was excluded from all pink noise analyses to avoid adaptation artifacts at sweep offset.

***spe-1: Simultaneous Patch and Neuropixels Dataset***
Simultaneous in vivo patch-clamp and Neuropixels recordings were obtained from Lister-Hooded rats under urethane anesthesia (1.4–1.8 g/kg i.p.). The dataset was provided by Marques-Smith et al. (2020), which describes the full data acquisition protocol in detail. Briefly, neurons were recorded in primary motor cortex and primary somatosensory cortex (forelimb, hindlimb, and trunk areas). Patch-clamp signals were acquired at ~50,023 Hz. The Neuropixels probe provided simultaneous LFP at 2,500 Hz and AP-band signals at 30,000 Hz.

Of 43 recorded neurons, 40 had sufficient spikes for the waveform clustering and LFP analyses; three cells (c17, c18, c43) were excluded due to insufficient patch-clamp spikes. Neuron c39 had no LFP data, and was therefore excluded from all LFP-based analyses, though it was included in the waveform-only clustering analysis. All 43 neurons were included in the intra-spike features correlation analyses (**Supp. Fig. 1**). All cell metadata described below (Supp. Fig. **6**) was assigned by Marques-Smith et al. Neurons were classified as pyramidal cells (PC, n = 35) or interneurons (IN, n = 5; cells c7, c12, c16, c18, c22) based on the trough-to-peak duration and half-width of the patch spike-triggered average (PSTA) of the highest-amplitude Neuropixels channel. Recording modality was juxtacellular for 35 neurons (27 in voltage-clamp, 8 in current-clamp configuration) and whole-cell patch-clamp for 5 neurons. Cortical depth ranged from 389 to 1,754 µm (median 1,146 µm), with the majority of neurons located between

700–1,500 μm, corresponding to Layer 5. Extracellular action potential (EAP) detectability was assessed via PSTA of the Neuropixels AP-band signal: 20 neurons had a clear EAP (peak-to-peak amplitude >10 μV), 5 were classified as dark neuron candidates (no detectable PSTA despite proximity to the probe in a high-yield session), and 15 lacked a detectable EAP attributable to probable technical factors.

***Preprocessing:*** Raw binary recordings were loaded and filtered using 4th-order Butterworth filters applied with zero-phase forward-backward filtering (SOS implementation). The patch-clamp signal was bandpass filtered at 10–25,000 Hz. The Neuropixels AP-band signal was bandpass filtered at 300–14,000 Hz. The LFP signal (Neuropixels LFP channel, 2,500 Hz) was used without additional software filtering, as it was already hardware-bandlimited by the probe acquisition system; because gain calibration was not available for this dataset, LFP amplitude is reported in raw, uncalibrated signal units rather than calibrated microvolts. Spikes were detected from the filtered patch-clamp trace as threshold crossings, with each cell's threshold set individually (range: 0.81–137.2 mV across cells, reflecting the range of recording modalities and signal amplitudes). Waveforms were extracted in a 5 ms pre- and 5 ms post-peak window. Following feature extraction, spikes with an inflection time outside valid bounds (physiologically implausible values indicating a fitting failure) were excluded.

### STATISTICAL REPORTING CONVENTIONS

Throughout, asterisks denote p-value magnitude (*$p < 0.05$, **$p < 0.01$, ***$p < 0.001$). Where a result is described as FDR corrected, the reported p-value and star tier reflect the Benjamini-Hochberg-corrected p-value, not the raw/uncorrected value.

### AP WAVEFORM PARAMETERIZATION (FIg. 2, Supp. Fig. 1-3)

We developed a novel three-part ramp-peak-decay parameterization approach (implemented in the open-source *spikewise* package; https://github.com/voytekresearch/spikewise) that quantifies fine-scale features of AP waveforms using an overcomplete set of intra-spike and inter-spike features. The parameterization pipeline is described in detail below.

***Control point detection.*** The spike derivative was smoothed using LOWESS (*statsmodels.nonparametric.lowess*; smoothing fraction = 0.008 for pvc-6, 0.01 for spe-1). The inflection point – the onset of the pre-spike ramp – was located as the intersection of two linear fits to the smoothed derivative: one over the -4 to -1 ms pre-peak window (slow ramp phase) and one from the point at which the derivative falls below its pre-peak standard deviation on the rising flank to the derivative peak (fast-rise phase). The ramp segment spans the voltage trace for 0.5 ms (pvc-6) or 1 ms (spe-1) immediately preceding the inflection point. Half-amplitude crossings flanking the peak were defined using the midpoint between inflection and peak voltages. The exponential window began 0.5 ms after the peak and extended 5 ms.

***Feature extraction.*** Nine features were extracted per spike. From the ramp: slope of a linear fit over the ramp segment (ramp_amp), time from inflection to peak (inflection_time), and voltage at the ramp end (inflection_amp). From the peak: peak voltage (peak_amp), duration between the half-amplitude crossings (peak_width), and mean voltage drop from the peak to ±0.1 ms around it (peak_sharpness). From the decay: rate (exp_lambda) and asymptotic offset (exp_const) of a three-parameter exponential fit $V(t) = A \cdot \exp(-\lambda t) + C$. Goodness of fit was assessed as the squared Pearson correlation between observed and fitted values ($R^2$). The $\log_{10}$ inter-spike interval (log_isi) was computed from spike times; the last spike per recording was assigned NaN.

***Configurable parameters*****.** Detection threshold, window length, smoothing fraction, ramp window (pre_peak_ms), ramp segment length (pre_inflection_ms), and exponential window onset and duration are all user-configurable in *spikewise.* Defaults were used throughout except where noted above (per-cell detection thresholds in spe-1; smoothing fraction and ramp segment length differing between datasets).

***Quality control.*** By default, spikes with an inflection time outside valid bounds are excluded, as this indicates a failed ramp/inflection fit; ±inf feature values are also set to NaN. spikewise additionally supports user-defined minimum $R^2$ thresholds (computed as squared Pearson correlation between observed and fitted values) for both the ramp and exponential decay fits, enabling stricter quality filtering if required. Remaining NaN values propagated through downstream analyses via NaN-aware operations.

***Recording stability (Supp. Fig. 3).*** To verify that waveform parameter variability did not increase over recording time (which would indicate patch degradation or a dying neuron) two analyses were applied to spe-1. First, the IQR was computed within 10 equal-width normalized-time bins per cell×feature, then normalized by that cell's mean IQR; a flat profile at 1.0 indicates stable variability. Second, for each cell×feature pair, the absolute deviation of each spike's feature value from the cell mean (normalized by the population IQR) was regressed against normalized recording time using linear regression; a positive slope would indicate increasing variability over time. Slopes were pooled across all cell×feature pairs and tested with a one-sided Wilcoxon signed-rank test ($H_1$: slope > 0).

### COMPUTING POWER SPECTRA AND SPECTRAL PARAMETERIZATION

**pvc-6.** For each pink noise spike, the power spectral density (PSD) was estimated over a window of injected current immediately preceding the spike's inflection point (200 ms for main results; sensitivity to window length is explored in supplementary analyses) using a single-segment Welch method (Hann window; NeuroDSP compute_spectrum; Cole et al., 2019). The DC component was discarded. The spectral exponent was computed as the negative slope of a linear fit to log10(power) versus log10(frequency).

**spe-1.** Three LFP spectral analyses were performed, each using different windowing but sharing the same spectral parameterization settings (Donoghue et al., 2020): aperiodic_mode = "fixed"; peak_width_limits = (4, 8) Hz; max_n_peaks = 4; peak_threshold = 1.5 SD; fit range 1–90 Hz. The aperiodic exponent and offset were extracted from each fit. Band-specific oscillatory power in theta (4–15 Hz) and gamma (30–80 Hz) was quantified as the area under the periodic component above the aperiodic fit (log-space, clipped at zero).

***Peri-transition analysis.*** LFP recordings were segmented into temporal blocks defined by detected waveform cluster transition boundaries. Welch PSDs were estimated per block (Hann window; nperseg = min(block length, 4 s); 50% overlap; scipy.signal.welch) and SpectralModel was fit to each block.

***LFP by cluster-transition occurrence.*** Welch PSDs (same parameters) were estimated over the full LFP recording per cell, yielding one spectrum per cell. Cells were then grouped by which spike waveform feature exhibited temporal cluster transitions.

***Per-spike spectral analysis (ridge regression)*****.** For each spike, a 2.5 s LFP window (1.0 s pre, 1.5 s post) was extracted. Multitaper PSDs were computed in sliding 0.5 s windows (4 ms step; tfr_array_multitaper; time_bandwidth = 2.0; 256 frequencies, 1–90 Hz) and SpectralTimeModel was fit to each time bin. Spectral features were averaged over two peri-spike windows: pre-spike (−55 to −5 ms) and post-spike (+5 to +55 ms).

**PVC-6: STIMULUS TYPE DECODING (Fig. 3A-C, Supp. Fig. 4)**

To test whether spike waveform features carry information about the type of injected current, three classifiers were trained to predict stimulus type (constant, ramp, or pink noise) from waveform features alone (8 features; log ISI and stimulus statistics excluded). Data were split 80/20 into training and test sets. Three decoders were evaluated: (1) multinomial logistic regression (lbfgs solver); (2) linear SVM with C selected by 5-fold cross-validated grid search ($C \in \{0.1, 1, 10, 100\}$); and (3) random forest with hyperparameters selected by 5-fold grid search (n_estimators ∈ {50, 100, 200}; max_depth ∈ {None, 10, 20}; min_samples_split ∈ {2, 5, 10}; min_samples_leaf ∈ {1, 2, 4}). Classification accuracy was estimated by bootstrapping ($n = 1{,}000$) the training set with replacement and evaluating on the held-out test set each iteration. Feature importances were averaged across bootstrap iterations to estimate mean importance and variability per feature.

For the random forest classifier specifically, the confusion matrix and feature importances reported in Fig. 3B-C use out-of-fold predictions: cross_val_predict-equivalent 5-fold cross-validation (same hyperparameters as above; class_weight unweighted) was repeated 1,000 times with independent random 5-fold partitions, yielding a stable accuracy estimate (mean ± SD) and an averaged confusion matrix; feature importances were pooled across all fold-level estimators from the same repeats. In this analysis, constant-current spikes were frequently misclassified as pink noise (Fig. 3B). This was more likely for sweeps with lower current, closer to the current level typically used for pink noise (Pearson $r = -0.78$, $p < 0.0001$, $n = 20$ sweeps), pointing to the importance of current amplitude in shaping AP waveform, consistent with the pink-noise-specific analysis.

**PVC-6: PINK NOISE STIMULUS REGRESSION (Fig. 3D-G, Supp. Fig. 5)**

Ridge regression was used to predict instantaneous pre-spike stimulus properties (spectral exponent, mean amplitude, and standard deviation) from all spike waveform features in pink noise trials. The final spike of each sweep was excluded to avoid edge effects at sweep boundaries. Both predictors and targets were z-scored prior to fitting (StandardScaler), so beta weights reflect SD change in the target per SD change in each waveform feature. The regularization parameter $\alpha$ was selected via RidgeCV ($\alpha \in$ logspace(−3, 3, 100)) on z-scored features, and model performance was evaluated by 5-fold cross-validation (shuffle = True). Model significance was assessed via permutation test ($n = 1{,}000$ label permutations). $R^2$ and 95% CIs were estimated by bootstrapping ($n = 1{,}000$). P-values were corrected for multiple comparisons across the three stimulus targets using false discovery rate (FDR) correction (Benjamini–Hochberg, $\alpha = 0.05$). Pairwise Pearson correlations between spike features and stimulus targets (Fig. 3E), and one-sample t-tests on bootstrapped beta weights against zero (Fig. 3G), were each FDR corrected (Benjamini-Hochberg, $\alpha = 0.05$) across all spike-feature × stimulus-target pairs tested (24 pairs each).

***Pre-spike window sensitivity.*** To assess how far back the stimulus history influences waveform shape, ridge regression was repeated across pre-inflection window sizes of 5, 25, 50, 100, 200, 300, 400, and 500 ms. Paired shuffle controls (permuted independently at each window size) establish the null $R^2$ at each window. Main results report the 200 ms window; beta weights and spike-stimulus correlations are additionally shown for selected windows.

**SPE-1: WITHIN-NEURON SPIKE FEATURE CLUSTER CHARACTERIZATION (Fig. 4B-C, Supp. Fig. 7-10)**

For each cell, spike feature distributions were examined to identify discrete groups of spikes – referred to as clusters – when a feature showed multimodal structure. To detect these cluster groups, spike counts were binned using data-adaptive bin widths (Freedman-Diaconis rule: $2 \times IQR \times n^{-1/3}$) and the histogram was smoothed with a triangular kernel ([1, 2, 3, 2, 1]). A peak in the smoothed histogram was counted as a distinct cluster if it reached at least 10% of the tallest peak height and the valley between adjacent peaks was at least 70% of the shorter flanking peak. Features with fewer than 8 unique values were skipped and solutions with more than 3 clusters were rejected. Each spike was assigned to its nearest peak and clusters were labeled Low, Mid, or High in ascending order of cluster mean (**Supp. Fig. 7**).

Where clusters were identified, waveform separation between cluster states was quantified using two metrics computed on mean cluster waveforms. 1) nRMSE (normalized RMSE) was computed after scaling both waveforms by their global amplitude maximum, such that it reflects differences in relative waveform amplitude between clusters while controlling for overall scale. 2) Cosine similarity was computed directly on the mean waveforms and captures shape differences independent of amplitude: values near 1 indicate near-identical waveform shape across clusters, lower values indicate greater shape divergence.

Waveform cluster assignment was examined in relation to inter-spike interval (ISI) to characterize whether spikes from different waveform cluster groups differed in firing rate. For each cell with both a waveform cluster and an ISI cluster, the association between waveform cluster membership and log ISI was quantified using $\eta^2$ (Kruskal-Wallis $H / (n - 1)$), which measures the proportion of rank-variance in log ISI explained by waveform cluster assignment. Values near 0 indicate independence. Across all cells with both cluster types ($n = 33$), the median $\eta^2$ was 0.013, indicating that waveform cluster membership does not systematically co-vary with firing rate (**Supp. Fig.** 9).

To assess whether cluster membership changed over the course of the recording, Spearman $\rho$ was computed between each spike's ordinal position in the recording (spike 1, 2, 3, ...) and its ordinal cluster label (Low = 0, Mid = 1, High = 2). A high positive $\rho$ indicates spikes progressively shift toward higher-valued clusters over time; a high negative $\rho$ indicates the reverse; values near zero indicate no temporal structure. Temporal-structure p-values were FDR corrected (Benjamini-Hochberg) across all cell × feature pairs (**Supp. Fig. 10C**). Associations with cell-level metadata were tested using Mann-Whitney U (binary variables), Kruskal-Wallis (multi-level), or Spearman $\rho$ (continuous), with FDR correction applied (**Supp. Fig. 10**).

**SPE-1: CLUSTER TEMPORAL TRAJECTORIES AND TRANSITION DETECTION (Supp. Fig. 11)**

Within each recording, we aimed to identify neurons whose AP waveform showed a clear shift between cluster states over time, and to pinpoint when that shift occurred. To identify these transitions, the ordinal cluster label sequence (Low = 0, Mid = 1, High = 2) was smoothed using a centered 50-spike rolling mean. One to three logistic sigmoids were then fit to the smoothed trajectory:

$$y = b + \Sigma_i L_i / (1 + \exp(-k_i(t - t_{0i})))$$

where $t_0$ is the transition time, $k$ is the transition rate (steepness), and $L$ is the amplitude. The number of sigmoids was selected by AIC. A transition was retained only if the overall model $R^2 \geq 0.80$ and the sharpest sigmoid had $|k| \geq 0.02\ s^{-1}$. Each accepted sigmoid inflection point $t_0$ was recorded as a transition event, with the dominant cluster state before and after the transition identified from the surrounding spike labels. Cell-feature pairs with fewer than 20 valid spikes were excluded.

**SPE-1: PERI-TRANSITION LFP ANALYSIS (Supp. Fig. 12)**

For each detected waveform cluster transition, three non-overlapping LFP blocks were extracted relative to the transition time $t_0$: pre (120 s ending 30 s before $t_0$), peri (60 s centered on $t_0$), and post (120 s beginning 30 s after $t_0$). Blocks shorter than 30 s were excluded. LFP power spectra were computed and parameterized for the pre and post blocks as described in the Spectral Analysis section, yielding mean amplitude, standard deviation, aperiodic exponent, aperiodic offset, theta area under the curve (AUC; that is, spectral power in the theta band, adjusted for the baseline aperiodic activity), and gamma AUC per block.

To account for transitions in either direction, each transition was assigned a sign based on its directionality: sign = +1 for upward transitions (cluster label increases from pre to post) and sign = −1 for downward transitions (label decreases). For upward transitions, the pre block corresponded to the lower cluster state and the post block to the higher cluster state; for downward transitions this assignment was reversed. This ensured that comparisons were always between equivalent lower and higher waveform cluster states regardless of transition direction. LFP features were compared between lower and higher cluster states using a Wilcoxon signed-rank test (paired by transition event).

**SPE-1: LFP ANALYSIS BY FEATURE TRANSITION OCCURRENCE (Supp. Fig. 13)**

To test whether the LFP oscillatory environment is associated with which spike waveform feature shows a cluster transition, cells were grouped by transitioning feature (e.g., all cells with a peak amplitude transition in one group, all cells with a log ISI transition in another). Cells with transitions in more than one feature contributed to each relevant group. Cells with no detected transitions formed a control group. For each cell, a Welch PSD was computed over the full LFP recording and parameterized as described in the Spectral Analysis section, yielding mean amplitude, standard deviation, aperiodic exponent, aperiodic offset, theta AUC, and gamma AUC. For spectral visualization, between-cell amplitude differences were removed by subtracting each cell’s mean log-power and adding back the group mean. LFP features were compared across all groups using a Kruskal-Wallis test per feature.

**SPE-1: WITHIN- VS BETWEEN-NEURON WAVEFORM VARIABILITY (Fig. 4D, Supp. Fig. 14)**

To contextualize the degree of within-cell spike waveform variability relative to across-cell differences, two types of distances were computed. For waveform-level distances (nRMSE and cosine similarity), each spike was compared to its own cell’s mean waveform (within-cell) or to the mean waveform of every other cell (between-cell). Within-cell nRMSE was normalized by the peak amplitude of the cell's own mean waveform; between-cell nRMSE was normalized by the larger of the two cells’ peak amplitudes. Between-cell comparisons were additionally broken down by cell type (PC/PC, IN/IN) and recording modality (juxtacellular/juxtacellular, whole-cell/whole-cell).

For feature-level distances, within-cell variability was quantified as the absolute value of the difference between the spike value  and the cell mean per spike, pooled across cells. Between-cell distances were computed as the absolute value of the difference between the spike value and the other cell mean for every spike–cell pair across all cell combinations, pooled across all pairs. Finally, for each cell and each feature, the percentage of within-cell spikes whose distance exceeded the mean of the global between-cell distribution was computed, providing a per-cell measure of how much within-cell variability overlaps with the between-cell range.

### SPE-1: LFP RIDGE REGRESSION (Fig. 4E-G, Supp. Fig. 15-16)

To test whether spike waveform features covary with the peri-spike LFP state, we used ridge regression to predict five LFP features (mean amplitude, standard deviation, aperiodic exponent, gamma AUC, and theta AUC) from eight spike waveform features (ramp amplitude, inflection time, inflection amplitude, peak amplitude, peak width, peak sharpness, decay λ, decay constant). LFP features were extracted from pre-spike (−55 to −5 ms) and post-spike (+5 to +55 ms) windows as described in the Spectral Analysis section. A log ISI–only model was run in parallel as a control to assess how much LFP variance is explained by firing rate alone, independent of waveform shape.

Within each cell, both predictors and targets were z-scored. The regularization parameter α was selected via RidgeCV (5-fold cross-validation; $\alpha \in$ logspace(−3, 3, 100)). Model significance was assessed via permutation test (n = 1,000 label permutations); per-cell significance was FDR corrected (Benjamini-Hochberg, α = 0.05) across cells within each LFP target and predictor set. Pre-spike and post-spike β weights were highly correlated across cells (Spearman $\rho \geq 0.89$ for LFP amplitude and variability targets; **Supp. Fig. 15**) and were averaged prior to supplementary population analysis. Population-level model performance was assessed per LFP target using a one-sided Wilcoxon signed-rank test ($H_1$: CV $R^2 > 0$), FDR corrected (Benjamini-Hochberg, α = 0.05) across LFP targets within each predictor set. Population-level β significance was assessed using a one-sample t-test ($H_1$: mean $\beta \neq 0$, α = 0.05, uncorrected across features), restricted to LFP targets with FDR-significant CV $R^2$.

To check whether high $R^2$ values could simply reflect low variance in the LFP targets, raw target variance (before z-scoring) was examined across cells with usable LFP recordings. Target standard deviation was never near zero, with a minimum of 0.18. Variance also did not correlate with $R^2$ the way a variance artifact would predict: for all four targets, pre- and post-spike LFP amplitude and LFP standard deviation, the correlation between target std and $R^2$ was weak and negative, not positive (r = -0.207, p = 0.199; r = -0.276, p = 0.085; r = -0.072, p = 0.657; r = -0.042, p = 0.796).

### SPE-1: STIMULATION ARTIFACT CHECK (Supp. Fig. 17)

Although the spe-1 dataset is described as spontaneous activity recorded without experimenter-controlled stimulus, we cannot rule out that stimulation was delivered at some points during these recordings to drive spiking activity. To assess this possibility, we checked each cell's raw, unfiltered patch voltage trace for shifts in resting membrane potential: if stimulation were provided, this may be seen as a sustained shift in baseline membrane potential, which the bandpass filter used elsewhere in this manuscript would attenuate. For each cell, the raw trace was binned into 1 s windows and summarized by the median voltage per bin. PELT (Pruned Exact Linear Time) changepoint detection (ruptures package; L2 cost, minimum segment length 5 bins) was applied to this per-second baseline signal with a penalty of 3 log(n) times the signal variance, flagging every point at which the mean baseline level shifted rather than only the single largest jump.

To test whether these baseline shifts corresponded to shifts in spike waveform shape, each spike's cluster assignment (Low/Mid/High) for every clustered waveform feature was recoded as an ordinal value (0/1/2), averaged into the same 1 s bins as the baseline trace, and analyzed with the same PELT procedure. A waveform-cluster changepoint was considered to coincide with a baseline changepoint if the two fell within 1 minute of each other.

Baseline changepoints were found in 28 of 29 cells with a usable (non-degenerate) baseline trace, and PELT changepoints were found in at least one clustered waveform feature for 21 of these cells. The minimum distance from each of 112 waveform-cluster changepoints to its cell's nearest baseline changepoint was closer than expected under a null built by redrawing changepoint times uniformly at random within each cell's recording (median distance 1.00 min vs. 1.29 min null, 2,000 permutations, $p = 0.029$). Among the 22 baseline-shift cells with a clustered waveform feature, 19 (86%) also showed a coincident waveform-shape shift, consistent with brief experimenter stimulation in at least some of these cells. Even so, this does not appear to be the sole driver of the waveform variability reported elsewhere in this manuscript: of those 19 cells, 15 (79%) still showed more than one cluster present among spikes following the changepoint rather than switching cleanly to a single dominant cluster.

### DATA AND MATERIALS AVAILABILITY

All data analyzed in this manuscript are from publicly available sources. The pvc-6 dataset (in vitro whole-cell patch-clamp recordings from mouse visual cortex; Berg, 2014) is available through the Collaborative Research in Computational Neuroscience data sharing initiative (CRCNS; http://dx.doi.org/10.6080/K0H12ZXD). The spe-1 dataset (simultaneous patch-clamp and Neuropixels recordings from rat somatosensory cortex; Marques-Smith et al., 2018) is available through CRCNS (http://dx.doi.org/10.6080/K0J67F4T). All analysis code, including spike waveform parameterization, clustering, LFP analysis, and ridge regression pipelines, is publicly available at https://github.com/voytekresearch/spikewise
